# Neuroimaging profiling identifies distinct brain maturational subtypes of youth with mood and anxiety disorders

**DOI:** 10.1101/2022.08.30.505946

**Authors:** R Ge, R Sassi, LN Yatham, S Frangou

## Abstract

Mood and anxiety disorders typically begin in adolescence and have overlapping clinical features but marked inter-individual variation in clinical presentation. The use of multimodal neuroimaging data may offer novel insights into the underlying brain mechanisms. We applied Heterogeneity Through Discriminative Analysis (HYDRA) to measures of regional brain morphometry, neurite density, and intracortical myelination to identify subtypes of youth, aged 9-10 years, with mood and anxiety disorders (N=1931) compared to typically developing youth (N=2823). We identified three subtypes that were robust to permutation testing and sample composition. Subtype 1 evidenced a pattern of imbalanced cortical-subcortical maturation compared to the typically developing group, with subcortical regions lagging behind prefrontal cortical thinning and myelination and greater cortical surface expansion globally. Subtype 2 displayed a pattern of delayed cortical and subcortical maturation indicated by higher cortical thickness and subcortical volume and lower cortical surface area expansion and myelination compared to the typically developing group. Subtype 3 showed evidence of atypical brain maturation involving globally lower cortical thickness and surface coupled with higher myelination and neural density. Subtype 1 had superior cognitive function in contrast to the other two subtypes that underperformed compared to the typically developing group. Higher levels of parental psychopathology, family conflict, and social adversity were common to all subtypes, with subtype 3 having the highest burden of adverse exposures. These analyses comprehensively characterize pre-adolescent mood and anxiety disorders, the biopsychosocial context in which they arise, and lay the foundation for the examination of the longitudinal evolution of the subtypes identified as the study sample transitions through adolescence.

## Introduction

Mood and anxiety disorders collectively account for over 65% of the disease burden attributable to mental disorders in individuals aged 10–24 years [1]. Most of the disability is accounted for by the early onset of these disorders [2] and their persistence into adulthood [3, 4]. There is international consensus on the need to prioritize youth mental health initiatives aiming at prevention and early intervention [5–7] which importantly acknowledges the major challenge posed by the incomplete understanding of the underlying neurobiology [8].

The diagnostic boundaries of mood (i.e., major depression, bipolar disorders) and anxiety disorders, as currently defined [9, 10] are highly permeable due to overlapping psychopathology; negative affective states in particular are both shared and central clinical features of mood and anxiety disorders [11] including bipolar disorder where depressive symptoms are the dominant psychopathology [12]. The clinical overlap between mood and anxiety disorders is even more pronounced in youth as depressive and anxiety symptoms typically predate the onset of syndromal diagnoses [13, 14]. Similarly, research into the neurobiological correlates of mood and anxiety disorders has failed to produce disorder-specific biomarkers but has identified instead transdiagnostic neuroimaging and genetic features [15–17].

In response, research efforts have shifted towards new approaches to classification that use data-driven methods to identify subgroups of individuals with mood and anxiety disorders based on their shared biological properties. This approach has yielded promising findings in adults [17–19] and has only recently begun to be applied in youth. Kaczkurkin and colleagues (2020) [20] identified two neuroanatomical subtypes in youth (aged 8-23 years) with internalising disorders broadly corresponding to an “impaired” and a “spared” group. A similar spared-impaired pattern was reported by Fan and colleagues (2021) [21] who used data-driven methods to distinguish 9-10-year-olds with internalizing disorders into two subgroups with either high or low impulsivity and then contrasted their neuroanatomical profiles.

The present study leverages the resources of the Adolescent Brain Cognitive Development (ABCD) Study (https://abcdstudy.org/) which acquires comprehensive phenotypic information and high-quality multimodal neuroimaging data from the largest currently available sample of 9-10-year-olds [22]. Our working hypothesis is that participants with mood and anxiety disorders can be partitioned into data-driven biologically informative subtypes based on their neuroimaging features. The study design enriches prior literature in three distinct ways. First, we expanded the phenotypic spectrum to encompass the entire range of mood and anxiety disorders, including bipolar disorder. Bipolar disorder is typically excluded from the construct of “internalizing” disorders despite compelling evidence for the dominance of internalizing symptoms, which is even more pronounced in young people [14]. Second, we expanded the neuroimaging phenotypic space to include measures of neurite density [23] and myelination [24, 25] that could yield insights into the brain maturational processes that may differentiate potential subtypes. Third, both psychopathology and brain organization are influenced by the environment in which individuals live. Prior research has emphasized the importance of perinatal history, parental socioeconomic status and psychopathology, family and neighborhood environment [26–28]; measures pertaining to these domains were tested to identify potentially distinct associations with the neuroimaging-based subtypes. These analyses comprehensively characterize pre-adolescent mood and anxiety disorders, the biopsychosocial context in which they arise, and lay the foundation for the investigation of the longitudinal evolution of the maturational and clinical trajectories of these subtypes in follow-up studies of this sample.

## Methods

### Sample

De-identified data were accessed from the 2.0 and 3.0 data releases (https://data-archive.nimh.nih.gov/abcd) of the ABCD study. Participants at baseline (N=11878) were recruited using multi-stage probability sampling to ensure that the sample is nationally representative in terms of sex, race/ethnicity, socio-economic status, and urbanicity [29] (Supplementary Material S1). The coordinating center is the University of California, San Diego, which has oversight of the ethical approval of the ABCD protocols. At each site, parents provided written informed consent, and youth provided assent.

The current study sample comprises ABCD participants selected based on the completeness of diagnostic data from the Kiddie Schedule for Affective Disorders and Schizophrenia for School-Age Children (KSADS-5) [30, 31] and high-quality neuroimaging data across all modalities based on ABCD quality control criteria [32] (Supplementary Material S2, Supplementary Table S1). The flowchart of the selection process is presented in Supplementary Figure S1.

Based on conventional psychiatric labels the study sample included the following five groups: (i) 2823 typically developing participants that were free of any lifetime psychopathology [age: 119.13 (7.50) months, 52.60% female]; (ii) 178 participants diagnosed only with a depressive disorder [age: 119.08 (7.08) months, 50.00% female]; (iii) 307 participants diagnosed only with a bipolar disorder [age: 118.65 (7.13) months, 51.79% female]; (iv) 1197 participants diagnosed only with an anxiety disorder [age: 118.97 (7.60) months, 55.89% female], and (v) 249 participants with comorbid mood and anxiety disorders [age: 119.00 (7.55) months, 54.62% female] (additional details in Supplementary Tables S2). Depressive disorders comprised major depressive disorder, dysthymia, or unspecified depressive disorder. Bipolar disorders comprised bipolar disorder I, bipolar disorder II, or unspecified bipolar disorder. Anxiety disorders comprised separation anxiety disorder, social anxiety disorder, generalized anxiety disorder, specific phobia, or other unspecified anxiety disorders. Further details of the diagnostic composition of the sample are provided in Supplementary Table S3.

### Non-imaging Measures

Participants’ internalizing and externalizing behaviors were assessed with the Child Behavior Checklist (CBCL) [33] and their crystallized and fluid intelligence were evaluated using the NIH Toolbox [34]. The PhenX ToolKit [35] and study-specific questionnaires [36–38] were used to record information about participants’ prenatal and obstetric history, parental characteristics (i.e., parental socioeconomic status and psychopathology), exposure to adverse life events, and quality of family, school and neighborhood environment (i.e., family function, quality of parental engagement, school performance and engagement and neighborhood deprivation). Definitions of these measures are provided in Supplementary Material S4 and Supplementary Table S4).

### Neuroimaging

The protocols for data acquisition, processing, and neuroimaging feature extraction [32,39] are outlined in Supplementary Material S5. Neuroimaging measures were extracted from whole-brain T_1_-weighted, and diffusion-weighted magnetic resonance imaging scans mapped Desikan-Killiany cortical parcellations [40] and subcortical regions mapped to the Fischl probabilistic atlas [41] and were accessed through the ABCD repository. These comprised (a) FreeSurfer derived measures of cortical thickness and surface area and regional subcortical volume; (b) Restriction Spectrum Imaging (RSI) [23] derived measures of neural density (ND); and (c) regional cortical myelination obtained through the gray/white matter contrast (GWC), which is the ratio of the signal intensity on either side of the surface formed at the gray-white matter cortical boundary [24, 25]. All neuroimaging measures were site harmonized using the ComBat algorithm (https://github.com/Jfortin1/ComBatHarmonization) [42] before further analyses.

### Statistical Analysis

#### Missing Data

Only participants with complete demographic, clinical, and neuroimaging data were included. Missingness for all other non-imaging measures was low (range 0.04%-8.50%) and missing values were imputed using multivariate imputation by chained equations (implemented in R 4.0.5 using the “mice” function) [43].

#### Semi-supervised Machine Learning for Subtype Identification

We employed Heterogeneity Through Discriminative Analysis (HYDRA), a nonlinear semi-supervised machine learning algorithm [44] to identify homogeneous clusters (i.e., subtypes) amongst pre-adolescents with mood and anxiety disorders. HYDRA over other clustering methods does not rely on similarity indices that are susceptible to the effects of non-specific factors such as age and sex but employs indices of deviation between a group of interest (here participants with mood and anxiety disorders) and a reference group (here typically developing participants) based on a combination of multiple hyperplanes (using a convex polytope) thus accommodating non-linearity (Supplementary Material S6; Supplementary Figure S2).

HYDRA (https://github.com/evarol/HYDRA) was applied to the regional morphometric, GWC, and ND measures. The diagnostic labels of the clinical group were not included in the input features of HYDRA while age, sex, and race/ethnicity were included as covariates. We assessed solutions with 2-7 clusters and chose the optimal solution following a 5-fold cross-validation based on the Adjusted Rand Index (ARI) [45]. Statistical significance was determined through permutation at P<0.05 (details in Supplementary Material S6).

#### Comparison of personal, family, and social characteristics of the identified subtypes

We compared each HYDRA cluster’s neuroimaging, personal, family, and social characteristics to the typically developing group using a series of mixed linear models implemented with the “lme4” package in R. The effects of age, sex, and race/ethnicity were regressed from all models and total intracranial volume (TIV) from models for cortical surface area, subcortical volume, and neurite density (Supplementary Figures S3 and S4). The threshold for statistical significance for these post-hoc analyses was set at P_FDR_<0.005 with false discovery rate (FDR) [46] correction for multiple testing.

### Sensitivity Analyses

We conducted two sensitivity analyses to test the robustness of the clustering solution to reproducibility and relatedness (details in Supplementary Material S7). We also tested the sensitivity of the results to alternate definitions of intracortical myelin by using the T1w/T2w [47] instead of GWC (details in Supplementary Material S7).

## Results

### HYDRA-Subtypes

The 3-cluster HYDRA solution had the highest ARI and was robust to permutation testing value (Supplementary Material S8; Supplementary Figure S5) and sample composition (Supplementary Material S9; Supplementary Figure S7). Each of these clusters (henceforth referred to as subtypes) comprised 631, 649, and 651 ABCD participants. The diagnostic composition of each subtype was generally proportional to the diagnostic distribution in the entire clinical sample (Figure 1); subtype 3 however had relatively fewer participants with anxiety disorders and relatively more with bipolar disorders compared to subtype 1 (χ^2^□=□15.80, P□=□0.001) and subtype 2 (χ^2^□=□10.12, P□=□0.02).

**Figure 1.**
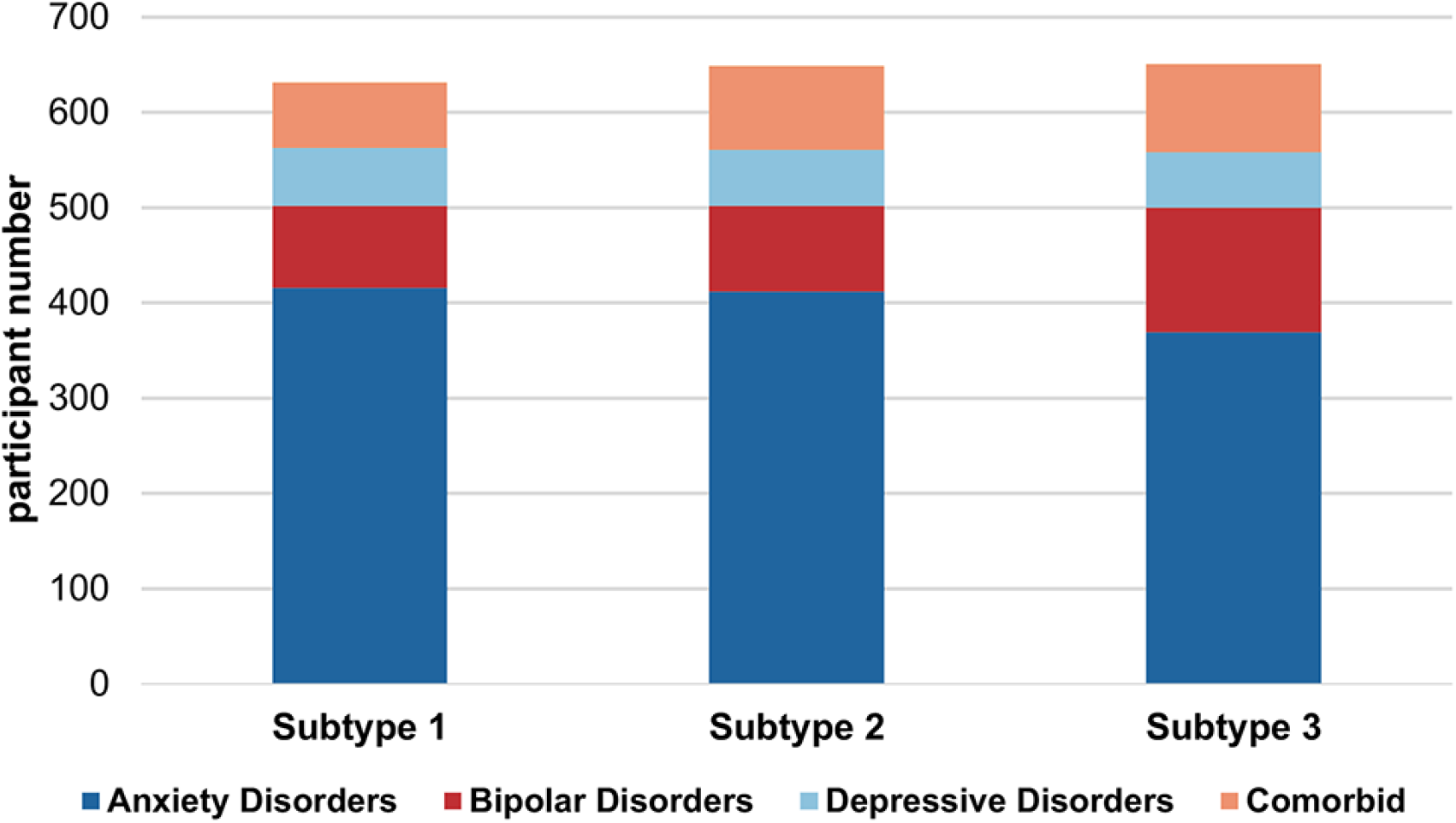
Transdiagnostic Composition of the HYDRA-identified Subtypes of Mood and Anxiety Disorders in Pre-adolescents. The figure illustrates the composition of each cluster in terms of traditional clinical diagnoses. Chi-squared tests revealed that subtype 1 (N=631) and subtype 2 (N=649) had equal proportions of participants with mood and anxiety disorders (χ^2^□=□2.45, P□=□0.48), and subtype 3 (N=651) had more participants with bipolar disorders, and fewer participants with anxiety disorders relative to subtype 1 (χ^2^□=□15.80, P□=□0.001) and subtype 2 (χ^2^□=□10.12, P□=□0.02).

### Clinical, Family, and Social Characteristics of the HYDRA-subtypes

The results of each HYDRA-subtype’s comparison to the typically developing group are summarized below and in Table 1 and Supplementary Table S5.

**Table 1.** Clinical, Family, and Social Characteristics of the HYDRA-Subtypes and the Typically Developing Group.

|  | **Typically**<br>**Developing**<br>**N=2823** | **Subtype 1**<br>**N=631** | **Subtype 2**<br>**N=649** | **Subtype 3**<br>**N=651** |
| --- | --- | --- | --- | --- |
| **Age** (months), mean (SD) | 119.13 (7.5) | 119.49 (7.42) | 118.1 (7.38) | 119.23 (7.55) |
| **Sex** (%) (male/female) | 47/53 | 47/53 | 43/57 | 48/52 |
| **Pubertal Stage** (%) (pre/early/mid/late/post-puberty) | 50/24/24/2/0 | 49/22/27/2/0 | 47/25/26/2/0 | 45/20/33/2/0 |
| **Race/Ethnicity** (%) (White/Black/Hispanic/Asian/Other) | 52/14/22/3/09 | 57/8/23/1/11 | 53/14/20/2/11 | 49/22/21/1/7 |
| **Obstetric and Perinatal** |  |  |  |  |
| **Birth weight**, (lbs) mean (SD) | 7 (1.47) | 7.26 (1.53)* | 7.04 (1.47) | 6.87 (1.47) |
| **Number of substances used by mother during pregnancy**, mean (SD) | 0.06 (0.29) | 0.09 (0.36)* | 0.09 (0.31)* | 0.11 (0.37)* |
| **Psychopathology and Cognition** |  |  |  |  |
| **CBCL-Internalizing**, mean (SD) | 43.21 (8.39) | 49.09 (9.59)* | 48.66 (9.47) * | 47.77 (9.46) * |
| **CBCL-Externalizing**, mean (SD) | 40.67 (7.45) | 43.62 (8.42) * | 44.16 (8.53) * | 43.76 (8.29) * |
| **Fluid Intelligence**, mean (SD) | 97.38 (17.3) | 97.38 (16.47) | 95.89 (16.84)* | 95.57 (16.29)* |
| **Crystallized Intelligence**, mean (SD) | 106.64 (18.26) | 110.09 (18.47)* | 105.06 (17.88)* | 103.87 (17.91)* |
| **Academic Performance and School Environment** |  |  |  |  |
| **Drop in Grade** (%) (yes/no) | 6/94 | 8/92 | 7/93 | 11/87* |
| **School Disengagement**, mean (SD) | 3.6 (1.36) | 3.69 (1.37) | 3.68 (1.47) | 3.76 (1.55)* |
| **School Involvement**, mean (SD) | 13.38 (2.13) | 13.16 (2.28)* | 13.1 (2.35)* | 13.1 (2.31)* |
| **Positive View of School**, mean (SD) | 20.31 (2.56) | 19.86 (2.64)* | 19.86 (2.73)* | 19.99 (2.93)* |
| **Parental Characteristics** |  |  |  |  |
| **Parental Cohabiting**, (%) (yes/no) | 78/22 | 77/23 | 76/24 | 71/29* |
| **Parents with Mental Health Problems**, (%) (both parents/one parent/neither parent) | 5/30/65 | 8/35/57* | 10/36/54* | 8/36/56* |
| **Parents with Help Seeking for Mental Problems**, (%) (both parents/one parent/neither parent) | 8/22/70 | 13/29/58* | 12/29/59* | 11/24/65* |
| **Family and Peer Environment** |  |  |  |  |
| **Family Conflict-Parental Report**, mean (SD) | 2.12 (1.76) | 2.32 (1.91)* | 2.36 (1.83)* | 2.24 (1.84)* |
| **Family Conflict – Child Report**, mean (SD) | 1.72 (1.76) | 1.91 (1.83)* | 2.06 (1.89)* | 2 (1.92)* |
| **Parental Engagement**, mean (SD) | 4.46 (0.46) | 4.43 (0.48) | 4.40 (0.50)* | 4.38 (0.51)* |
| **Regular Group of Friends, (%) (yes/no)** | 92/8 | 93/7 | 91/9 | 88/12* |
| **Experienced Bullying, (%) (yes/no)** | 6/94 | 13/87* | 13/87* | 14/86* |
| **Exposure to Stressful Events, (%) (yes/no)** | 27/73 | 34/66* | 34/66* | 34/66* |
| **Socioeconomic Environment** |  |  |  |  |
| **Parental Education, mean (SD)** | 16.64 (2.8) | 16.91 (2.51)* | 16.51 (2.96) | 16.27 (2.72)* |
| **Family Income Level, mean (SD)** | 7.32 (2.43) | 7.48 (2.21)* | 7.1 (2.45)* | 6.77 (2.49)* |
| **Financial Adversity, mean (SD)** | 0.27 (0.83) | 0.4 (0.98)* | 0.41 (0.95)* | 0.52 (1.13)* |
| **Area Deprivation Index, mean (SD)** | 37.48 (26.22) | 37.73 (26.47) | 39.35 (27.91) | 44.69 (27.98)* |
| SD=standard deviation; %=percentage; Definitions of all the variables are provided in Supplementary Table S4. * P<0.005, FDR corrected; between each subtype and typically developing group. |  |  |  |  |

#### Obstetric and Perinatal events

Maternal use of alcohol, substances, and prescribed medications was higher in all subtypes while subtype 1 had higher birth weight.

#### Psychopathology and Cognition

All subtypes had higher CBCL scores for externalizing and internalizing psychopathology; subtype 1 had higher crystallized intelligence while subtype 3 had lower crystallized and fluid intelligence.

#### Academic Performance and School Environment

Participants in all subtypes reported less positive school involvement and had a less positive view of their school environment as a place of opportunity and support but only participants in subtype 3 reported higher alienation from academic goals and were more likely to drop in grades

#### Parental characteristics

The prevalence of parental psychiatric diagnosis and helpseeking for mental health problems were higher in all subtypes while the parents of participants in subtype 3 were less likely to live together.

#### Family and Peer Environment

Greater family conflict and higher exposure to adverse life events and bullying were common to all subtypes. Additionally, participants in subtypes 2 and 3 reported less parental engagement and those in subtype 3 were also less likely to have a regular group of friends.

#### Socioeconomic environment

ABCD participants in subtype 1 came from families with higher family income and higher education level, but participants in all subtypes experienced periods of financial adversity. Only ABCD participants in subtype 3 lived in neighborhoods with a higher area deprivation index.

### Neuroimaging Profiles of the HYDRA-subtypes

Figure 2 presents the differences between each subtype and the typically developing group for each neuroimaging phenotype at P_FDR_<0.005. Subtype 1 was comparable to the typically developing group in terms of neurite density but had larger amygdala, striatal, and thalamic volumes and globally greater surface area expansion; they also had lower cortical thickness and higher intra-cortical myelin in dorsal prefrontal regions. Subtype 2 evidenced globally higher cortical thickness and globally lower cortical surface expansion and cortical myelin, as well as lower neurite density in ventromedial prefrontal, parietal, and occipital cortices. Subtype 3 was characterized by lower cortical thickness primarily in posterior brain regions and globally lower surface area, lower subcortical volume coupled with higher cortical myelination and neural density.

**Figure 2.**
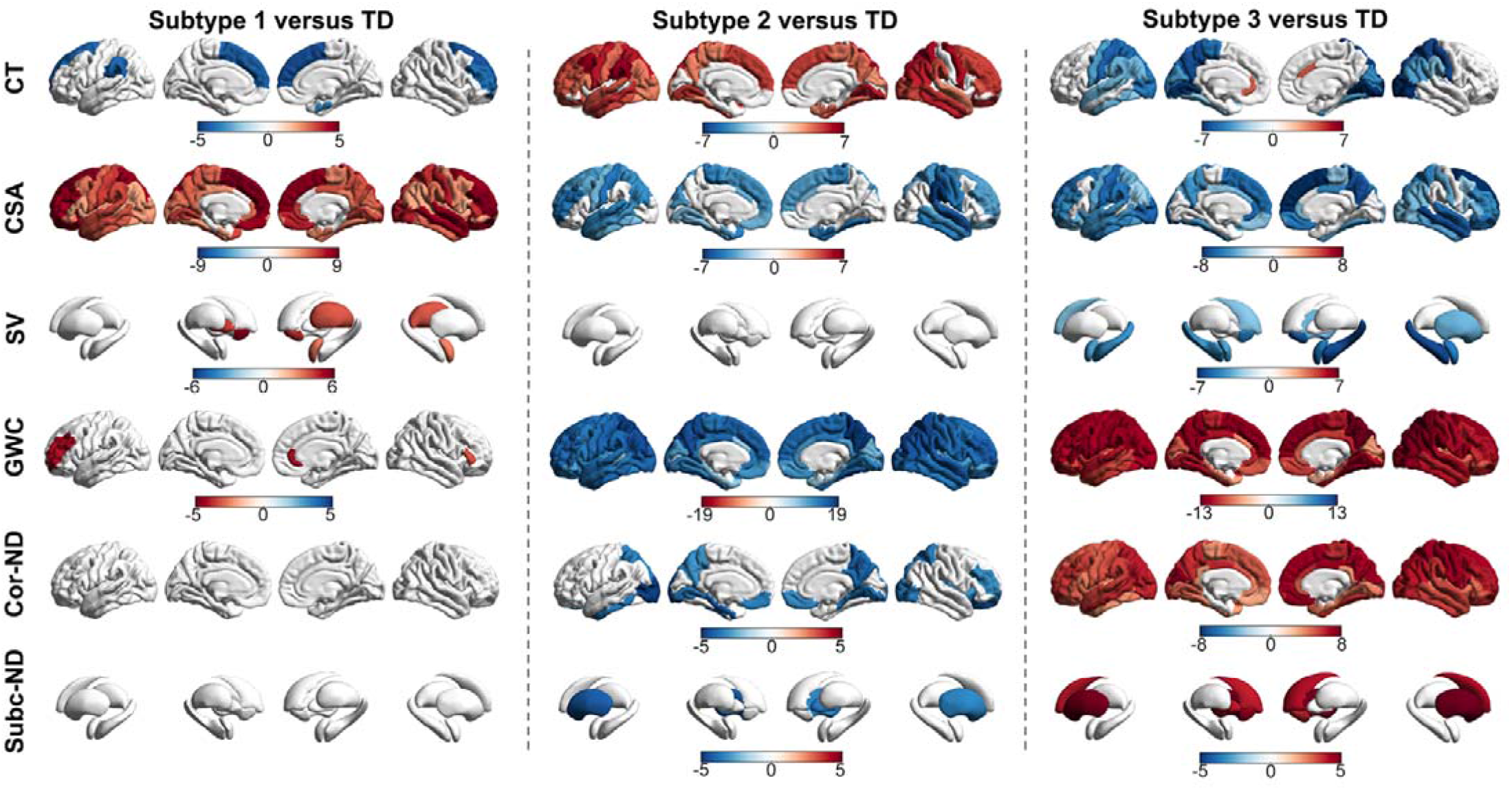
Neuroimaging Profiles of the HYDRA-identified Subtypes of Mood and Anxiety Disorders in Pre-adolescents. Regional differences (represented by the corresponding t-values) between each subtype and the typically developing (TD) group in cortical thickness (CT), cortical surface area (CSA), subcortical volumes (SV), neurite density (ND) and gray/white matter contrast (GWC). Only significant differences at P_FDR_<0.005 are shown. Warm and cool colors respectively indicate higher and lower values in each subtype compared to the TD group; however, for visualization purposes, in the GWC plots warm and cool colors respectively indicate lower and higher values in each subtype compared to the TD group.

## Discussion

We leveraged the power of machine learning with the unique resources of the ABCD study to test for the presence of neuroimaging-defined subtypes in juvenile mood and anxiety disorders. We identified three transdiagnostic subtypes that evidenced greater exposure to parental psychopathology, family conflict, and adverse experiences including bullying, compared to the typically developing group. Beyond these similarities, the three subtypes had distinct neuroimaging profiles potentially implicating different developmental mechanisms.

### The HYDRA subtypes transcend clinical diagnoses

The subtypes identified using neuroimaging-based clustering included a mixture of ABCD participants with mood and anxiety disorders that may reflect diagnostic equivocality with some youth showing diagnostic continuity while others may change diagnostic status or may recover [48–50]. However, neuroimaging studies in adults with established and chronic diagnoses of mood and anxiety disorders have also failed to identify diagnosis-specific changes in neuroimaging phenotypes suggesting that the mechanisms underlying symptom expression transcend diagnostic categories regardless of age [15, 17].

### The HYDRA subtypes had similar exposures to adversity

Risk factors for mood and anxiety disorders have been widely researched. The literature provides robust evidence that risk factors are not diagnosis-specific but are shared across multiple disorders. These include familial predisposition [51–53], perinatal exposure to substances and medications [54, 55], multiple indicators of lower socioeconomic status (lower parental education, low family income, and neighborhood deprivation) [56, 57], family dysfunction, and poor parental practices [58, 59], unfavorable view of the school environment [60] and exposure to a range of adverse experiences, including difficulties in forming peer-relationships and bullying [61]. The findings of the current study largely reinforce this pattern with the exception of subtype 1, in which several socioeconomic indicators relating to family income and education and positive parental engagement were better than those in the typically developing group. Subtype 3 evidence a greater burden of adversity with greater negative exposures across most domains.

### The HYDRA subtypes had distinct neuroimaging profiles

Prior neuroimaging studies in mood and anxiety disorders in youth have focused exclusively on brain morphometry using diagnosis-specific case-control comparisons. Excessive cortical thinning and lower surface area have been reported in youth depression [62, 63] and mania/hypomania [64]. Some studies have associated juvenile anxiety disorders with both higher and lower cortical thickness in different prefrontal areas, and lower subcortical volumes [65–67] but others found no casecontrol differences [68, 69]. A key supposition of these neuroimaging studies was that each diagnostic group would be associated with a distinct pattern of brain structural alterations underpinning the specific disorder-related symptoms. The present findings do not support such formulation as the three subtypes identified were transdiagnostic.

Instead, our results suggest that different brain maturational profiles can give rise to a range of clinical symptoms. Typical brain maturational events that dominate late childhood and adolescence are cortical myelination along a posterior-to-anterior gradient [70] and adaptive synaptic/dendritic density reduction [71, 72]. Associated brain morphometric changes involve cortical thinning [25, 73, 74], cortical surface area expansion [75], and a gradual reduction in subcortical volumes from childhood onwards [76, 77]. During typical development, cortical thinning, surface area expansion, and cortical myelination have been associated with higher cognitive abilities [78–81], Conversely, excessive cortical thinning and diminished cortical surface area expansion have been associated with lower cognitive abilities [82, 83].

In this context, the neuroimaging profile of subtype 1 suggests a pattern of imbalanced cortical-subcortical maturation compared to the typically developing group, with subcortical regions lagging behind prefrontal cortical thinning and myelination and greater cortical surface expansion globally. This interpretation is aligned with cortical maturation being associated with better cognitive abilities [78–81] which were also characteristic of subtype 1 (Figure 2, Table 1). The neuroimaging profile of subtype 2 is suggestive of a pattern of generally delayed cortical and subcortical maturation evidenced by higher cortical thickness and subcortical volume and lower cortical surface area expansion and myelination compared to the typically developing group (Figure 2). This pattern was also associated with cognitive disadvantage (Table 1). Subtype 3 showed evidence of atypical brain maturation involving globally lower cortical thickness and surface coupled with higher myelination and neural density and significant disadvantage in cognition, academic performance, and peer relationships (Figure 2; Table 1). These findings can be interpreted as potential evidence of abnormalities in cell size and packing density which resonates with abnormalities in brain maturation seen in neurodevelopmental disorders [84, 85].

The ABCD study includes planned follow-up assessments using the same protocols that cover a 10-year period of development and will thus enable us to assess the value of neuroimaging-based classification of mood and anxiety disorders identified here. Specifically, the availability of longitudinal data will enable the comparison of this approach to traditional diagnosis-based models for the prediction of functional outcomes of ABCD participants, with regards to cognition, psychopathology and social functioning.

### Methodological Considerations

Compared to previous studies, the large sample size, narrow age range, and the epidemiological approach to recruitment are major strengths as they optimize the capacity to examine brain variability in mood and anxiety disorders while minimizing challenges arising from selection bias and the inclusion of participants at different developmental stages. The use of HYDRA is a methodological advance over other clustering techniques as it enables the identification of clinical subgroups that differ from the normative reference group along multiple dimensions. Analyses of data from subsequent follow-up waves of the ABCD study will enable the investigation of the longitudinal evolution of the clusters identified and their prognostic values for mental health.

Taken together, the findings of this study uncovered brain structural heterogeneity in youth with mood and anxiety disorders indicating marked differences in maturational profiles but lack diagnostic specificity. Future follow-up analyses will enhance understanding of the functional consequences for cognition and psychopathology of further brain development as ABCD participants transition through adolescence.

## Supplementary information is available at MP’s website

## Acknowledgements

N/A

## Conflict of Interest

RG, RS and SF report no conflicts of interest. LNY is a consultant and/or has received speaker fees and/or sits on the advisory board and/or receives research funding from Abbvie, Alkermes, Allergan, Canadian Network for Mood and Anxiety Treatments (CANMAT), Canadian Institutes of Health Research (CIHR), Dainippon Sumitomo Pharma, Gedeon Richter, GSK, Intracellular Therapies, Lundbeck, Merck, Otsuka, Sanofi and Sunovion over the past 3 years.

