## Supplement for "Neuroimaging profiling identifies distinct brain maturational subtypes of youth with mood and anxiety disorders": Supplement.docx

**Supplementary Material**

**Supplementary Methods**

**S1. Overview of the ABCD study**

**S2. Study sample selection**

Supplementary Table S1. Diagnostic codes and variable names from the Kiddie Schedule for Affective Disorders and Schizophrenia for School-Age Children

Supplementary Figure S1. Flowchart of study sample selection

**S3. Study Sample Characteristics**

Supplementary Table S2. Study sample characteristics

Supplementary Table S3. Detailed diagnostic composition of youth with mood and anxiety disorders

**S4. Definition of the non-imaging variables**

Supplementary Table S4. Definition of non-imaging variables

**S5. Neuroimaging acquisition, processing, and feature extraction**

**S6. Heterogeneity Through Discriminative Analysis (HYDRA)**

Supplementary Figure S2. Schematic representation of HYDRA

Supplementary Figure S3. Correlations between total intracranial volume (TIV) with global measures of cortical thickness, cortical surface area, and subcortical volume

Supplementary Figure S4. Correlations between total intracranial volume (TIV) and global gray/white matter contrast and neurite density

**S7. Sensitivity Analyses**

Reproducibility

Intracortical Myelin

Relatedness

**Supplementary Results**

**S8. HYDRA-subtypes of mood and anxiety disorders**

Supplementary Figure S5. Adjusted Rand Index for HYDRA Models

Supplementary Table S5. Clinical, family, and social characteristics of HYDRA-subtypes

**S9. Sensitivity Analyses**

Reproducibility

Intracortical Myelin

Supplementary Figure S6. Cortical maps for GWC and T1/T2 ratio within each HYDRA subtype and the typically developing group

Relatedness

Supplementary Figure S7. Neuroimaging Profiles of the HYDRA-identified Subtypesin the entire sample and after exclusion of all related individuals

**References**

**S1. The ABCD Study**

The ABCD study is a prospective population-based multisite study that aims to follow the physical, mental, cognitive, and brain maturation trajectories of 9–10-year-olds for up to 10 years. All caregivers provided written informed consent and all children provided assent. The coordinating centre is the University of California, San Diego, which oversaw the ethical approval of the ABCD protocols. Prior to participation, parents or guardians provided written informed consent, and the children provided assent.

The recruitment catchment areas of the participating sites encompass over 20% of the entire US population 9–10-year-olds. The study used multi-stage probability sampling to ensure that the ABCD sample matched, as closely as possible, the sociodemographic variation of the US population [1]. The sociodemographic selection factors were age, sex, race and ethnicity, socio-economic status, and urbanicity. The stages involved were: (a) selection of a nationally distributed set of primary study sites, (b) probability sampling of schools within the defined catchment area of each site, and (c) recruitment of eligible children in each sample school.

**S2. Study Sample Selection**

First, we excluded participants with poor brain scan quality and low post-processing scores (based on the ABCD quality control criteria). Participants were further excluded if they had any of the following diagnoses: alcohol and substance use disorders, attention deficit hyperactivity disorder, conduct disorder, eating disorders, obsessive-compulsive disorder, oppositional defiant disorder, autism spectrum or psychotic spectrum disorders or trauma-related and stress disorders (Supplementary Table S1). Psychiatric diagnoses were established using the Kiddie Schedule for Affective Disorders and Schizophrenia for School-Age Children (KSADS-5) [2, 3]. These exclusions also applied to participants with mood and anxiety disorders. Of the remaining participants, those with self or parentally reported mood and anxiety disorders comprised the clinical population while participants with no psychiatric diagnosis comprised the typically developing group (Supplementary Figure S1). The issue of participant relatedness was addressed in the sensitivity analyses (Supplementary Material S7).

| **Table S1. Diagnostic coding and variables based on the Kiddie Schedule for Affective Disorders and Schizophrenia for School-Age Children (KSADS-5) and Schizophrenia for School-Age Children (KSADS-5)** | | | |
| --- | --- | --- | --- |
| **Disorder** | **Diagnostic Code** | **Variable name**  **child report** | **Variable name parental report** |
| **anxiety disorders** | panic disorder (F41.0) PRESENT | ksads_5_857_t | ksads_5_857_p |
|  | panic disorder (F41.0) PAST | ksads_5_858_t | ksads_5_858_p |
|  | other specified anxiety disorder (panic disorder impairment does not meet full criteria) F41.8 | ksads_5_906_t | ksads_5_906_p |
|  | diagnosis - other specified anxiety disorder PAST (panic disorder impairment does not meet full criteria) F41.8 | ksads_5_907_t | ksads_5_907_p |
|  | agoraphobia (F40.00) PRESENT | ksads_6_859_t | ksads_6_859_p |
|  | agoraphobia (F40.00) PAST | ksads_6_860_t | ksads_6_860_p |
|  | other specified anxiety disorder (Agoraphobia impairment does not meet full criteria) F41.8 | ksads_6_908_t | ksads_6_908_p |
|  | separation anxiety disorder (F93.00) PRESENT | ksads_7_861_t | ksads_7_861_p |
|  | separation anxiety disorder (F93.00) PAST | ksads_7_862_t | ksads_7_862_p |
|  | other specified anxiety disorder (separation anxiety disorder impairment does not meet full criteria) F41.8 | ksads_7_909_t | ksads_7_909_p |
|  | other specified anxiety disorder (separation anxiety disorder impairment does not meet full criteria) PAST F41.8 | ksads_7_910_t | ksads_7_910_p |
|  | social anxiety disorder (F40.10) PRESENT | ksads_8_863_t | ksads_8_863_p |
|  | social anxiety disorder (F40.10) PAST | ksads_8_864_t | ksads_8_864_p |
|  | other specified anxiety disorder (Social Anxiety Disorder impairment does not meet minimum duration) F41.8 | ksads_8_911_t | ksads_8_911_p |
|  | other specified anxiety disorder (social anxiety disorder impairment does not meet minimum duration) PAST F41.8 | ksads_8_912_t | ksads_8_912_p |
|  | specific phobia PRESENT (F40.2XX) | ksads_9_867_t | ksads_9_867_p |
|  | specific phobia PAST (F40.2XX) | ksads_9_868_t | ksads_9_868_p |
|  | generalized anxiety disorder Present (F41.1) | ksads_10_869_t | ksads_10_869_p |
|  | generalized anxiety disorder Past (F41.1) | ksads_10_870_t | ksads_10_870_p |
|  | other specified anxiety disorder (generalized anxiety disorder impairment does not meet minimum duration) F41.8 | ksads_10_913_t | ksads_10_913_p |
|  | other specified anxiety disorder (generalized anxiety disorder impairment does not meet minimum duration) PAST F41.8 | ksads_10_914_t | ksads_10_914_p |
|  | selective mutism (F94.0) PRESENT | ksads_25_865_t | ksads_25_865_p |
|  | selective mutism (F94.0) PAST | ksads_25_866_t | ksads_25_866_p |
|  | other specified anxiety disorder (selective mutism does not meet minimum duration) F41.8 | ksads_25_915_t | ksads_25_915_p |
|  | other specified anxiety disorder (selective mutism does not meet minimum duration) PAST F41.8 | ksads_25_916_t | ksads_25_916_p |
| **(2) bipolar disorders** | bipolar I disorder current episode manic (F31.1x) | ksads_2_830_t | ksads_2_830_p |
|  | bipolar I disorder current episode depressed F31.3x | ksads_2_831_t | ksads_2_831_p |
|  | bipolar I disorder currently hypomanic F31.0 | ksads_2_832_t | ksads_2_832_p |
|  | bipolar I disorder most recent Past episode manic (F31.1x) | ksads_2_833_t | ksads_2_833_p |
|  | bipolar I disorder most recent Past episode depressed (F31.1.3x) | ksads_2_834_t | ksads_2_834_p |
|  | bipolar II disorder currently hypomanic F31.81 | ksads_2_835_t | ksads_2_835_p |
|  | bipolar II disorder currently depressed F31.81 | ksads_2_836_t | ksads_2_836_p |
|  | bipolar II disorder most recent Past hypomanic F31.81 | ksads_2_837_t | ksads_2_837_p |
|  | unspecified bipolar and related disorder current (F31.9) | ksads_2_838_t | ksads_2_838_p |
|  | unspecified bipolar and related disorder PAST (F31.9) | ksads_2_839_t | ksads_2_839_p |
| **(3) depressive disorders** | major depressive disorder Present | ksads_1_840_t | ksads_1_840_p |
|  | major depressive disorder current in partial remission (F32.4) | ksads_1_841_t | ksads_1_841_p |
|  | major depressive disorder Past (F32.9) | ksads_1_842_t | ksads_1_842_p |
|  | persistent depressive disorder (dysthymia) PRESENT F34.1 | ksads_1_843_t | ksads_1_843_p |
|  | persistent depressive disorder (dysthymia) in partial remission F34.1 | ksads_1_844_t | ksads_1_844_p |
|  | persistent depressive disorder (dysthymia) PAST F34.1 | ksads_1_845_t | ksads_1_845_p |
|  | unspecified depressive disorder current (F32.9) | ksads_1_846_t | ksads_1_846_p |
|  | unspecified depressive disorder PAST (F32.9) | ksads_1_847_t | ksads_1_847_p |
| **(4) alcohol-related disorders** | alcohol use disorder Present | ksads_19_891_t | ksads_19_891_p |
|  | alcohol use disorder Past | ksads_19_892_t | ksads_19_892_p |
|  | unspecified alcohol-related disorder Present (F10.99) | ksads_19_893_t | ksads_19_893_p |
|  | unspecified alcohol-related disorder Past (F10.99) | ksads_19_894_t | ksads_19_894_p |
| **(5) attention-deficit/hyperactivity disorder** | attention-deficit/hyperactivity disorder Present | ksads_14_853_t | ksads_14_853_p |
|  | attention-deficit/hyperactivity disorder Past | ksads_14_854_t | ksads_14_854_p |
|  | attention-deficit/hyperactivity disorder in partial remission | ksads_14_855_t | ksads_14_855_p |
|  | unspecified attention-deficit/hyperactivity disorder (F90.9) | ksads_14_856_t | ksads_14_856_p |
| **(6) conduct disorder** | conduct disorder Present childhood onset (F91.1) | ksads_16_897_t | ksads_16_897_p |
|  | conduct disorder Present adolescent onset (F91.2) | ksads_16_898_t | ksads_16_898_p |
|  | conduct disorder Past childhood onset (F91.1) | ksads_16_899_t | ksads_16_899_p |
|  | conduct disorder Past adolescent onset (F91.2) | ksads_16_900_t | ksads_16_900_p |
| **(7) eating disorders** | anorexia nervosa (F50.02) binge eating/purging subtype PRESENT | ksads_13_929_t | ksads_13_929_p |
|  | anorexia nervosa (F50.02) binge eating/purging subtype present in partial remission | ksads_13_930_t | ksads_13_930_p |
|  | anorexia nervosa (F50.02) binge eating/purging subtype PAST | ksads_13_931_t | ksads_13_931_p |
|  | anorexia nervosa (F50.01) restricting subtype PRESENT | ksads_13_932_t | ksads_13_932_p |
|  | anorexia nervosa (F50.01) restricting subtype present in partial remission | ksads_13_933_t | ksads_13_933_p |
|  | anorexia nervosa (F50.01) restricting subtype PAST | ksads_13_934_t | ksads_13_934_p |
|  | bulimia nervosa (F50.2) PRESENT | ksads_13_935_t | ksads_13_935_p |
|  | bulimia nervosa (F50.2) PAST | ksads_13_936_t | ksads_13_936_p |
|  | bulimia nervosa partial remission Present | ksads_13_937_t | ksads_13_937_p |
|  | binge-eating disorder (F50.8) CURRENT | ksads_13_938_t | ksads_13_938_p |
|  | binge-eating disorder (F50.8) current in partial remission | ksads_13_939_t | ksads_13_939_p |
|  | binge-eating disorder (F50.8) PAST | ksads_13_940_t | ksads_13_940_p |
|  | other specified feeding or eating disorder anorexia nervosa current does not meet full criteria (F50.8) | ksads_13_941_t | ksads_13_941_p |
|  | other specified feeding or eating disorder bulimia nervosa current does not meet full criteria (F50.8) | ksads_13_942_t | ksads_13_942_p |
|  | other specified feeding or eating disorder bulimia nervosa PAST does not meet full criteria (F50.8) | ksads_13_943_t | ksads_13_943_p |
|  | other specified feeding or eating disorder binge eating disorder Present does not meet full criteria (F50.8) | ksads_13_944_t | ksads_13_944_p |
| **(8) obsessive-compulsive disorder** | obsessive-compulsive disorder Present (F42) | ksads_11_917_t | ksads_11_917_p |
|  | obsessive-compulsive disorder Past (F42) | ksads_11_918_t | ksads_11_918_p |
|  | other specified obsessive-compulsive and related disorders Present does not meet full criteria (F42) | ksads_11_919_t | ksads_11_919_p |
|  | other specified obsessive-compulsive and related disorders Past does not meet full criteria (F42) | ksads_11_920_t | ksads_11_920_p |
| **(9) oppositional defiant disorder** | oppositional defiant disorder Present F91.3 | ksads_15_901_t | ksads_15_901_p |
|  | oppositional defiant disorder Past F91.3 | ksads_15_902_t | ksads_15_902_p |
| **(10) schizophrenia spectrum and other psychotic disorders** | unspecified schizophrenia spectrum and other psychotic disorders F29 (Current) | ksads_4_851_t | ksads_4_851_p |
|  | unspecified schizophrenia spectrum and other psychotic disorders F29 (Past) | ksads_4_852_t | ksads_4_852_p |
| **(11) substance-related disorders** | cannabis use disorder Present | ksads_20_871_t | ksads_20_871_p |
|  | stimulant use disorder Present: amphetamine-type substance | ksads_20_872_t | ksads_20_872_p |
|  | sedative-hypnotic or anxiolytic use disorder Present | ksads_20_873_t | ksads_20_873_p |
|  | stimulant use disorder Present: cocaine | ksads_20_874_t | ksads_20_874_p |
|  | opioid use disorder Present | ksads_20_875_t | ksads_20_875_p |
|  | other hallucinogen use disorder Present | ksads_20_876_t | ksads_20_876_p |
|  | phencyclidine (PCP) use disorder: Present | ksads_20_877_t | ksads_20_877_p |
|  | inhalant use disorder Present | ksads_20_878_t | ksads_20_878_p |
|  | other drugs use disorder Present | ksads_20_879_t | ksads_20_879_p |
|  | cannabis use disorder Past | ksads_20_880_t | ksads_20_880_p |
|  | stimulant use disorder Past: amphetamine-type substance | ksads_20_881_t | ksads_20_881_p |
|  | Sedative-hypnotic or anxiolytic use disorder Past | ksads_20_882_t | ksads_20_882_p |
|  | stimulant use disorder Past: cocaine | ksads_20_883_t | ksads_20_883_p |
|  | opioid use disorder Past | ksads_20_884_t | ksads_20_884_p |
|  | other hallucinogen use disorder Past | ksads_20_885_t | ksads_20_885_p |
|  | phencyclidine (PCP) use disorder Past | ksads_20_886_t | ksads_20_886_p |
|  | inhalant use disorder Past | ksads_20_887_t | ksads_20_887_p |
|  | other drugs use disorder Past | ksads_20_888_t | ksads_20_888_p |
|  | substance use disorder CURRENT | ksads_20_889_t | ksads_20_889_p |
|  | substance use disorder PAST | ksads_20_890_t | ksads_20_890_p |
|  | unspecified substance-related disorder Present (F10.99) | ksads_20_893_t | ksads_20_893_p |
|  | unspecified substance-related disorder Past (F10.99) | ksads_20_894_t | ksads_20_894_p |
| **(12) trauma-and stressor-related disorders** | post-traumatic stress disorder PRESENT (F94.1) | ksads_21_921_t | ksads_21_921_p |
|  | post-traumatic stress disorder PAST (F94.1) | ksads_21_922_t | ksads_21_922_p |
|  | other specified trauma-and stressor-related disorder Present (PTSD impairment does not meet full criteria (F43.8) | ksads_21_923_t | ksads_21_923_p |
|  | other specified trauma-and stressor-related disorder PAST PTSD impairment does not meet full criteria (F43.8) | ksads_21_924_t | ksads_21_924_p |
| **(13) suicidal problems (used as an exclusion criterion for typically developing youth with no psychiatric diagnosis)** | self-injurious behavior without suicidal intent Present | ksads_23_945_t | ksads_23_945_p |
|  | suicidal ideation passive Present | ksads_23_946_t | ksads_23_946_p |
|  | suicidal ideation active nonspecific Present | ksads_23_947_t | ksads_23_947_p |
|  | suicidal ideation active method Present | ksads_23_948_t | ksads_23_948_p |
|  | suicidal ideation active intent Present | ksads_23_949_t | ksads_23_949_p |
|  | suicidal ideation active plan Present | ksads_23_950_t | ksads_23_950_p |
|  | preparatory actions toward imminent suicidal behavior Present | ksads_23_951_t | ksads_23_951_p |
|  | interrupted attempt Present | ksads_23_952_t | ksads_23_952_p |
|  | aborted attempt Present | ksads_23_953_t | ksads_23_953_p |
|  | suicide attempt Present | ksads_23_954_t | ksads_23_954_p |
|  | self-injurious behavior without suicidal intent Past | ksads_23_956_t | ksads_23_956_p |
|  | suicidal ideation passive Past | ksads_23_957_t | ksads_23_957_p |
|  | suicidal ideation active nonspecific Past | ksads_23_958_t | ksads_23_958_p |
|  | suicidal ideation active method Past | ksads_23_959_t | ksads_23_959_p |
|  | suicidal ideation active intent Past | ksads_23_960_t | ksads_23_960_p |
|  | suicidal ideation active plan Past | ksads_23_961_t | ksads_23_961_p |
|  | preparatory actions toward imminent suicidal behavior Past | ksads_23_962_t | ksads_23_962_p |
|  | interrupted attempt Past | ksads_23_963_t | ksads_23_963_p |
|  | aborted attempt Past | ksads_23_964_t | ksads_23_964_p |
|  | suicide attempt Past | ksads_23_965_t | ksads_23_965_p |

.

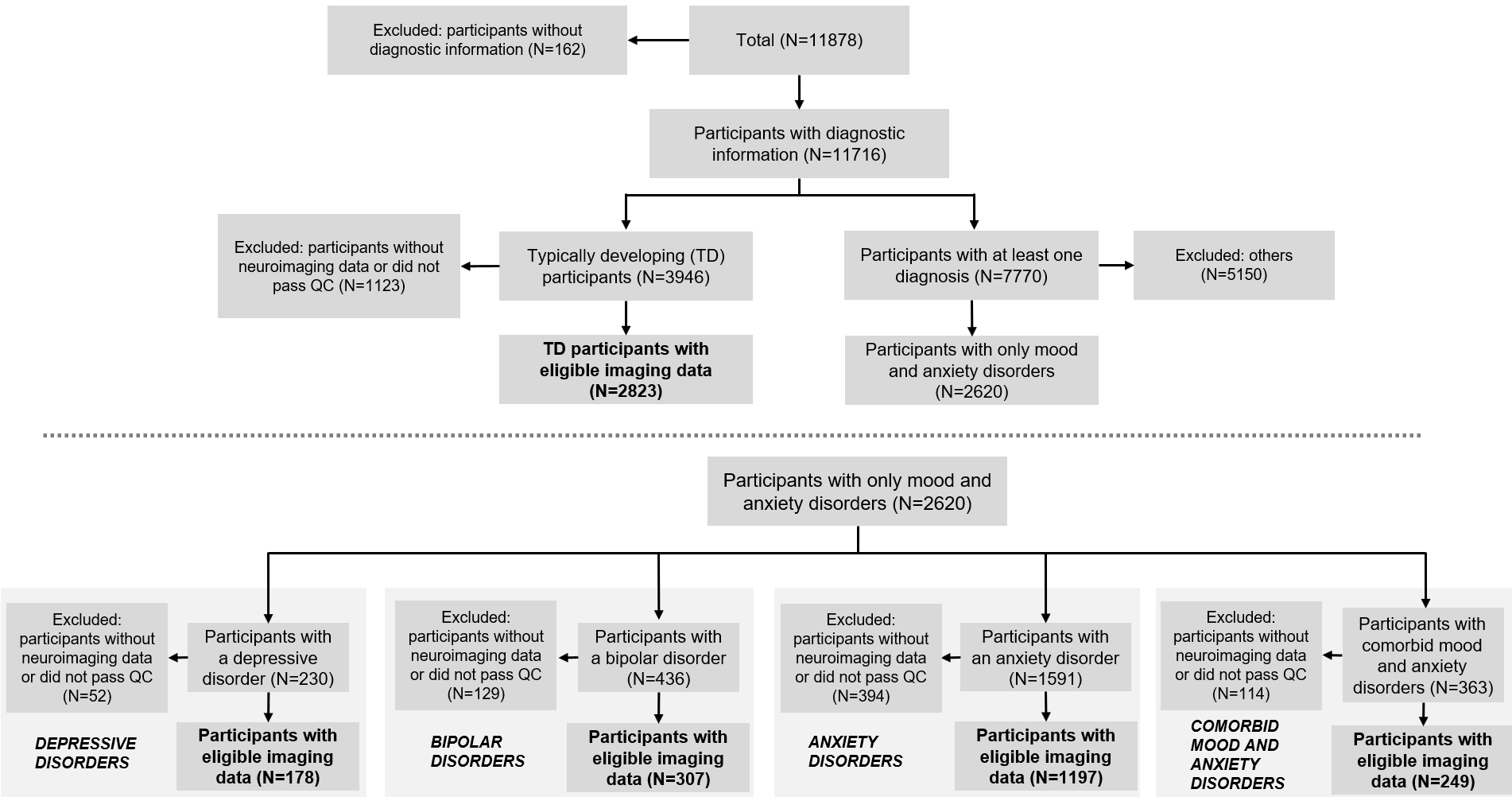

**Supplementary Figure S1. Flowchart of study sample selection**

**S3. Study Sample Characteristics**

| Supplementary Table S2. Study sample characteristics | | | | | |
| --- | --- | --- | --- | --- | --- |
|  | **Typically**  **Developing**  **N=2823** | **Depressive Disorders**  **N=178** | **Bipolar**  **Disorders**  **N=307** | **Anxiety**  **Disorders**  **N=1,197** | **Comorbid Mood and Anxiety Disorders**  **N=249** |
| Demographic Characteristics |  |  |  |  |  |
| Age (months), mean (SD) | 119.13 (7.5) | 119.08 (7.08) | 118.65 (7.13) | 118.97 (7.6) | 119 (7.55) |
| Sex (%male/%female) | 47/53 | 50/50 | 48/52 | 44/56 | 45/55 |
| Pubertal Stage (%pre/%early%/mid/%late/%post puberty) | 50/24/24/2/0 | 49/17/33/1/0 | 47/24/27/2/0 | 47/24/28/2/0 | 43/20/34/2/1 |
| Race/Ethnicity (%White/%Black/%Hispanic/%Asian/%Other) | 52/14/22/3/09 | 47/17/22/2/12 | 44/24/21/12/9 | 57/12/21/1/9 | 47/18/22/1/12 |
| Pregnancy and Birth | | | | | |
| Number of Obstetric Complications, mean (SD) | 0.32 (0.68) | 0.34 (0.71) | 0.37 (0.75) | 0.36 (0.76) | 0.39 (0.72) |
| Birth Weight (pounds), mean (SD) | 7 (1.47) | 7.04 (1.53) | 6.89 (1.52) | 7.09 (1.46) | 7.1 (1.6) |
| Premature Birth (%yes/%no) | 19/81 | 16/84 | 19/81 | 19/81 | 18/82 |
| Number of Maternal Medical Conditions During Pregnancy, mean (SD) | 0.85 (2.78) | 0.92 (2.82) | 0.98 (2.94) | 0.97 (2.88) | 1.55 (3.68) |
| Maternal Alcohol/Substance Use During Pregnancy | 0.06 (0.29) | 0.1 (0.43) | 0.11 (0.37) | 0.09 (0.32) | 0.12 (0.38) |
| Psychopathology and Cognition | | | | | |
| CBCL-Internalizing, mean (SD) | 43.21 (8.39) | 47.98 (9.95) | 44.63 (9) | 48.93 (9.12) | 51.56 (10.19) |
| CBCL-Externalizing, mean (SD) | 40.67 (7.45) | 44.41 (9.02) | 43.63 (8.81) | 43.47 (8.06) | 45.55 (8.93) |
| Fluid Intelligence, mean (SD) | 97.38 (17.3) | 93.12 (15.9) | 93.75 (16.26) | 97.7 (16.46) | 94.72 (17.04) |
| Crystallized Intelligence, mean (SD) | 106.64 (18.26) | 102.8 (17.53) | 102.4 (17.97) | 108.07 (18.25) | 105.11 (18.2) |
| Academic Performance and School Environment | | | | | |
| Drop in Grades (%yes/%no) | 6/94 | 12/88 | 10/90 | 7/93 | 13/87 |
| In-school Emotional Support (%yes/%no) | 0.2/99.8 | 0/1 | 0/1 | 0.3/99.7 | 1/99 |
| In-School Learning Support (%yes/%no) | 1/99 | 2/98 | 1/99 | 1/99 | 2/98 |
| In-School Aid (%yes/%no) | 1/99 | 0/1 | 2/98 | 1/99 | 1/99 |
| School Disengagement, mean (SD) | 3.6 (1.36) | 3.95 (1.7) | 3.81 (1.49) | 3.64 (1.39) | 3.78 (1.56) |
| School Involvement, mean (SD) | 13.38 (2.13) | 12.4 (2.8) | 13.2 (2.23) | 13.23 (2.21) | 13 (2.46) |
| Positive View of School, mean (SD) | 20.31 (2.56) | 19.08 (3.28) | 19.97 (2.8) | 20.03 (2.63) | 19.78 (2.9) |
| Parental Characteristics | | | | | |
| Parental Education | 16.64 (2.8) | 15.85 (2.67) | 16.11 (2.89) | 16.81 (2.64) | 16.41 (2.98) |
| Parental Employment (%yes/%no) | 72/28 | 63/37 | 69/31 | 70/30 | 92/28 |
| Parents in Partnership (%yes/%no) | 78/22 | 72/28 | 71/29 | 77/23 | 69/31 |
| Parental Psychiatric Diagnosis (%both parents/%one parent/%none) | 5/30/65 | 12/38/50 | 6/35/59 | 8/35/57 | 15/37/48 |
| Parental Help Seeking for Mental Problems (%both parents/%one parent/%none) | 8/22/70 | 10/27/63 | 06/27/67 | 13/27/60 | 13/32/55 |
| Family and Peer Environment | | | | | |
| Family Income | 7.32 (2.43) | 6.44 (2.48) | 6.67 (2.66) | 7.4 (2.23) | 6.8 (2.63) |
| Number of Indicators of Financial Adversity, mean (SD) | 0.27 (0.83) | 0.62 (1.24) | 0.45 (1) | 0.39 (0.97) | 0.53 (1.11) |
| Family Conflict-Parental Report, mean (SD) | 2.12 (1.76) | 2.4 (1.86) | 2.25 (1.86) | 2.27 (1.82) | 2.48 (2.03) |
| Family Conflict – Child Report, mean (SD) | 1.72 (1.76) | 2.72 (2.05) | 2.35 (1.95) | 1.71 (1.74) | 2.37 (2.04) |
| Primary Caregiver’s Warmth, mean (SD) | 2.81 (0.27) | 2.73 (0.33) | 2.74 (0.31) | 2.82 (0.26) | 2.78 (0.3) |
| Parental Supervision, mean (SD) | 4.46 (0.46) | 4.3 (0.59) | 4.37 (0.51) | 4.43 (0.48) | 4.36 (0.48) |
| Having Best Friend (%yes/%no) | 85/15 | 87/13 | 87/13 | 86/14 | 86/14 |
| Having a Regular Group of Friends (%yes/%no) | 92/8 | 85/15 | 88/12 | 93/7 | 87/13 |
| Bullying (%yes/%no) | 6/94 | 18/82 | 16/84 | 10/90 | 21/79 |
| Exposure to Stressful Events (yes/no) | 27/73 | 32/68 | 31/69 | 35/65 | 34/66 |
| Neighborhood Environment | | | | | |
| Area Deprivation Index, mean (SD) | 37.48 (26.22) | 46.33 (27.58) | 45.07 (30.06) | 37.81 (26.15) | 44.55 (29.65) |
| Continuous variables are shown as mean (standard deviation; SD) and categorical variables as percentage (%); Definitions of all the variables are provided in Supplemental Table S4. | | | | | |

| **Supplementary Table S3. Detailed diagnostic composition of youth with mood and anxiety disorders** | |
| --- | --- |
| **Depressive Disorders** | 62 (34.83%) have a history of major depressive disorder;  109 (61.24%) have a history of unspecified depressive disorder;  2 (1.12%) have a history of major depressive disorder and persistent depressive disorder;  5 (2.81%) have a history of major depressive disorder and unspecified depressive disorder. |
| **Bipolar Disorders** | 110 (35.83%) have a history of bipolar I disorder;  11 (3.58%) have a history of bipolar II disorder;  164 (53.42%) have a history of unspecified bipolar disorder;  9 (2.93%) have a history of bipolar I disorder and bipolar II disorder;  11 (3.58%) have a history of bipolar I disorder and unspecified bipolar disorder;  1 (0.33%) has a history of bipolar II disorder and unspecified bipolar disorder;  1 (0.33%) has a history of bipolar I disorder, bipolar II disorder, and unspecified bipolar disorder. |
| **Anxiety Disorders** | 5 (0.42%) have a history of agoraphobia;  74 (6.18%) have a history of separation anxiety disorder;  49 (4.09%) have a history of social anxiety disorder;  677 (56.56%) have a history of specific phobia;  20 (1.67%) have a history of generalized anxiety disorder;  142 (11.86%) have a history of other specified anxiety disorders;  1 (0.08%) has a history of panic disorder and other specified anxiety disorders;  1 (0.08%) has a history of agoraphobia and separation anxiety disorder;  1 (0.08%) has a history of agoraphobia and specific phobia;  1 (0.08%) has a history of agoraphobia and other specified anxiety disorder;  1 (0.08%) has a history of generalized anxiety disorder and other specified anxiety disorders;  8 (0.67%) have a history of separation anxiety disorder and social anxiety disorder;  62 (5.18%) have a history of separation anxiety disorder and specific phobia;  4 (0.33%) have a history of separation anxiety disorder and generalized anxiety disorder;  5 (0.42%) have a history of separation anxiety disorder and other specified anxiety disorder;  23 (1.92%) have a history of social anxiety disorder and specific phobia;  3 (0.25%) have a history of social anxiety disorder and generalized anxiety disorder;  7 (0.58%) have a history of social anxiety disorder and other specified anxiety disorder;  6 (0.50%) have a history of specific phobia and generalized anxiety disorder;  59 (4.93%) have a history of specific phobia and other specified anxiety disorder;  3 (0.25%) have a history of agoraphobia, specific phobia, and other specified anxiety disorder;  3 (0.25%) have a history of separation anxiety disorder, specific phobia, and generalized anxiety disorder;  4 (0.33%) have a history of separation anxiety disorder, specific phobia, and other specified anxiety disorder;  13 (1.09%) have a history of separation anxiety disorder, social anxiety disorder, and specific phobia;  2 (0.17%) have a history of separation anxiety disorder, social anxiety disorder, and other specified anxiety disorder;  9 (0.75%) have a history of social anxiety disorder, and specific phobia, and other specified anxiety disorder;  3 (0.25%) have a history of specific phobia, generalized anxiety disorder, and other specified anxiety disorder;  1 (0.08%) have a history of panic disorder, separation anxiety disorder, specific phobia, and other specified anxiety disorder;  1 (0.08%) have a history of agoraphobia, separation anxiety disorder, social anxiety disorder, and specific phobia;  3 (0.25%) have a history of separation anxiety disorder, social anxiety disorder, specific phobia, and generalized anxiety disorder;  1 (0.08%) have a history of separation anxiety disorder, social anxiety disorder, specific phobia, and other specified anxiety disorder;  3 (0.25%) have a history of social anxiety disorder, specific phobia, generalized anxiety disorder, and other specified anxiety disorder;  1 (0.08%) have a history of panic disorder, separation anxiety disorder, specific phobia, generalized anxiety disorder, and other specified anxiety disorder;  1 (0.08%) have a history of separation anxiety disorder, social anxiety disorder, specific phobia, generalized anxiety disorder, and other specified anxiety disorder. |
| **Comorbid Mood and Anxiety Disorders** | 3 (1.20%) have a history of separation anxiety disorder and unspecified bipolar disorder;  3 (1.20%) have a history of separation anxiety disorder and bipolar I disorder;  6 (2.41%) have a history of separation anxiety disorder and unspecified bipolar disorder;  6 (2.41%) have a history of separation anxiety disorder and unspecified depressive disorder;  28 (11.24%) have a history of specific phobia and bipolar I disorder;  1 (0.40%) have a history of specific phobia and bipolar II disorder;  33 (13.25%) have a history of specific phobia and unspecified bipolar disorder;  10 (4.02%) have a history of specific phobia and major depressive disorder;  24 (9.64%) have a history of specific phobia and unspecified depressive disorder;  3 (1.20%) have a history of generalized anxiety disorder and bipolar I disorder;  2 (0.80%) have a history of generalized anxiety disorder and major depressive disorder;  2 (0.80%) have a history of generalized anxiety disorder and unspecified depressive disorder;  13 (5.22%) have a history of other specified anxiety disorder and bipolar I disorder;  3 (1.20%) have a history of other specified anxiety disorder and bipolar II disorder;  15 (6.02%) have a history of other specified anxiety disorder and unspecified bipolar disorder;  7 (2.81%) have a history of other specified anxiety disorder and major depressive disorder;  9 (3.61%) have a history of other specified anxiety disorder and unspecified depressive disorder;  3 (1.20%) have a history of separation, specified anxiety disorder, and bipolar I disorder;  1 (0.40%) have a history of separation anxiety disorder, bipolar I disorder, and unspecified bipolar disorder;  3 (1.20%) have a history of separation, specific phobia, unspecified bipolar disorder;  1 (0.40%) have a history of separation, specific phobia, and unspecified depressive disorder;  1 (0.40%) have a history of separation, specific phobia, and major depressive disorder;  1 (0.40%) have a history of separation, other specified anxiety disorder, and unspecified depressive disorder;  2 (0.80%) have a history of social, specific phobia, and unspecified depressive disorder;  1 (0.40%) have a history of social, generalized anxiety disorder, and major depressive disorder;  1 (0.40%) have a history of social, generalized anxiety disorder, and unspecified depressive disorder;  2 (0.80%) have a history of social, generalized anxiety disorder, and unspecified bipolar disorder;  1 (0.40%) have a history of social, other specified anxiety, and major depressive disorder;  1 (0.40%) have a history of social, other specified anxiety, and unspecified depressive disorder;  3 (1.20%) have history of specific phobia, generalized anxiety, and unspecified bipolar disorder;  1 (0.40%) have a history of specific phobia, other specified anxiety disorder, and bipolar I disorder;  1 (0.40%) have a history of specific phobia, other specified anxiety disorder, and bipolar II disorder;  9 (3.61%) have a history of specific phobia, other specified anxiety disorder, and unspecified bipolar disorder;  4 (1.61%) have a history of specific phobia, other specified anxiety disorder, and major depressive disorder;  8 (3.21%) have a history of specific phobia, other specified anxiety disorder, and unspecified depressive disorder;  1 (0.40%) have a history of specific phobia, bipolar I disorder, and bipolar II disorder;  1 (0.40%) have a history of specific phobia, bipolar I disorder, and unspecified bipolar disorder;  1 (0.40%) has a history of specific phobia, major depressive disorder, and persistent depressive disorder;  1 (0.40%) have a history of generalized anxiety disorder, other specified anxiety disorder, and bipolar II disorder;  4 have history of other specified anxiety disorder, bipolar I disorder, and bipolar II disorder;  1 (0.40%) has history of other specified anxiety disorder, bipolar I disorder, and unspecified bipolar disorder;  4 (1.61%) have a history of other specified anxiety disorder, major depressive disorder, and unspecified depressive disorder;  1 (0.40%) have a history of separation, specific phobia, other specified anxiety, or major depressive disorder;  1 (0.40%) have a history of separation, specific phobia, other specified anxiety, unspecified depressive disorder;  1 (0.40%) have a history of separation, specific phobia, bipolar I disorder, unspecified bipolar disorder;  1 (0.40%) have a history of separation, generalized anxiety disorder, bipolar II disorder, unspecified bipolar disorder;  2 (0.80%) have history of separation, social, specific phobia, major depressive disorder;  1 (0.40%) have a history of separation, social, specific phobia, bipolar I disorder;  1 (0.40%) have a history of separation, social, other specified anxiety disorder, and major depressive disorder;  1 (0.40%) have a history of separation, social, generalized anxiety disorder, unspecified bipolar disorder;  3 (1.20%) have history of social, specific phobia, other specified anxiety disorder, and unspecified depressive disorder;  1 (0.40%) have a history of social, specific phobia, other specified anxiety disorder, and major depressive disorder;  1 (0.40%) have a history of social, specific phobia, major depressive disorder, and unspecified depressive disorder;  1 (0.40%) have a history of specific phobia, other specified anxiety disorder, major depressive disorder, and unspecified depressive disorder;  1 (0.40%) have a history of agoraphobia, separation anxiety disorder, specific phobia, other specified anxiety disorder, and unspecified depressive disorder;  2 (0.80%) have a history of separation anxiety disorder, social anxiety disorder, specific phobia, generalized anxiety disorder, and unspecified depressive disorder;  1 (0.40%) have a history of separation anxiety disorder, social anxiety disorder, specific phobia, generalized anxiety disorder, or major depressive disorder;  1 (0.40%) have a history of separation anxiety disorder, social anxiety disorder, specific phobia, generalized anxiety disorder, other specified anxiety disorder, and major depressive disorder. |
| Each participant may have more than one diagnosis | |

**S4. Definition of non-imaging variables**

Details of the procedures and instruments used in the ABCD study to assess the personal, family, and social characteristics of the participants have been previously published.2-4 They comprise study-specific questionnaires and widely used instruments such as the KSADS-5 [2,3], the Child Behavior Checklist (CBC) [4], and the PhenX7, and the Children's Report of Parental Behavioral Inventory [5], and the NIMH ToolBox [6]. A data dictionary providing detailed definitions of all the variables can be accessed at https://nda.nih.gov/data_dictionary.html. Below we provide details on the variables that we used in the analyses presented the current study.

| **Supplementary Table S4. Definition of non-neuroimaging measures** | | | |
| --- | --- | --- | --- |
| **Measure** | **Instrument** | **ABCD Variable Name** | **Coding** |
| Sex | ABCD Child Demographics | acspsw03: sex | Male or female; self-report |
| Age | ABCD Child Demographics | acspsw03: interview_age | Age in chronological months |
| Pubertal stage  Parent report | ABCD Parent Pubertal Development Scale and Menstrual Cycle Survey History (PDMS) | abcd_ssphp01:  pds_p_ss_female_category_2; pds_p_ss_male_category_2 | 1: Prepuberty; 2: Early puberty 3: Mid puberty; 4: Late puberty 5: Post puberty |
| Race/ethnicity | ABCD Child Demographics | acspsw03: race_ethnicity | 1 = White; 2 = Black; 3 = Hispanic; 4 = Asian; 5 = Other |
| **Obstetric and Perinatal History** | | | |
| Obstetric complications  Parent report | ABCD Developmental History Questionnaire; Participants answered the following question: “Did he/she have any of the following complications at birth? Blue at birth, Slow heartbeat, Did not breathe at first, Convulsions, Jaundice needing treatment, Required oxygen, Required blood transfusion, Rh incompatibility” | dhx01:  devhx_14a3_p, devhx_14b3_p devhx_14c3_p, devhx_14d3_p devhx_14e3_p, devhx_14f3_p devhx_14g3_p, devhx_14h3_p | Responses for each condition mentioned below were coded as:  1=present; 0=absent.  The sum of all responses was used.  devhx_14a3_p: Blue at birth  devhx_14b3_p: Slow heart  beatdevhx_14c3_p: Did not breathe at  firstdevhx_14d3_p: Convulsions  devhx_14e3_p: Jaundice needing treatment  devhx_14f3_p: Required oxygen  devhx_14g3_p: Required blood transfusion  devhx_14h3_p: Rh incompatibility  The sum of all the responses was used in the analyses |
| Birth weight  Parent report | ABCD Developmental History Questionnaire | dhx01: birth_weight_lbs | Birth weight in pounds |
| Premature birth  Parent report | ABCD Developmental History Questionnaire; Participants answered the following question: “Was the child born prematurely?” | dhx01: devhx_12a_p | 0=No; 1=Yes |
| Maternal medical conditions during pregnancy  Parent report | ABCD Developmental History Questionnaire; Participants answered the following question: “During the pregnancy with this child, did you/biological mother have any of the following conditions? Severe nausea and vomiting extending past the 6th month or accompanied by weight loss, Heavy bleeding requiring bed rest or special treatment, Pre-eclampsia, eclampsia, or toxemia, Severe gall bladder attack, Persistent proteinuria, Rubella (German measles) during first 3 months of pregnancy, Severe anemia, Urinary tract infections, Pregnancy-related diabetes, Pregnancy-related high blood pressure, Previa, Abruptio, or other problems with the placenta, An accident or injury requiring medical care, Any other conditions requiring medical care” | dhx01:  devhx_10a3_p, devhx_10b3_p devhx_10c3_p, devhx_10d3_p devhx_10e3_p, devhx_10f3_p devhx_10g3_p, devhx_10h3_p devhx_10i3_p, devhx_10j3_p devhx_10k3_p, devhx_10l3_p devhx_10m3_p | Responses for each condition below were coded as:  1=yes; 0=no.  The sum of all responses was used  devhx_10a3_p: Severe nausea and vomiting extending past the 6th month or accompanied by weight loss  devhx_10b3_p: Heavy bleeding requiring bed rest or special treatment  devhx_10c3_p: Pre-eclampsia, eclampsia, or toxemia  devhx_10d3_p: Severe gall bladder attack  devhx_10e3_p: Persistent proteinuria  devhx_10f3_p: Rubella (German measles) during first 3 months of pregnancy  devhx_10g3_p: Severe anemia  devhx_10h3_p: Urinary tract infections  devhx_10i3_p: Pregnancy-related diabetes  devhx_10j3_p: Pregnancy-related high blood pressure  devhx_10k3_p: Previa, abruptio, or other problems with the placenta  devhx_10l3_p: An accident or injury requiring medical care  devhx_10m3_p: Any other conditions requiring medical care  .The sum of the responses was used in the analyses |
| Maternal alcohol/substance use during pregnancy  Parent report | ABCD Developmental History Questionnaire; Participants answered the following question: “Once the biological mother knew she was pregnant, was she using any of the following? Tobacco, Alcohol, Marijuana, Cocaine, Heroin/Morphine, Oxycontin” | dhx01:  devhx_9_tobacco  devhx_9_alcohol devhx_9_marijuana devhx_9_coc_crack devhx_9_her_morph devhx_9_oxycont | Responses for each substance below were coded as:  1=used; 0=not used  The sum of all responses was used  devhx_9_tobacco: Tobacco  devhx_9_alcohol: Alcohol  devhx_9_marijuana: Marijuana  devhx_9_coc_crack: Cocaine/Crack  devhx_9_her_morph: Heroin/Morphine  devhx_9_oxycont: Oxycontin |
| **Psychopathology and Cognition** | | | |
| Internalizing Problems | Child Behavior Checklist (CBCL) | cbcl_scr_syn_internal_t | CBCL-Internal CBCL Syndrome Scale (t-score) |
| Externalizing Problems | Child Behavior Checklist (CBCL) | cbcl_scr_syn_external_t | CBCL-External CBCL Syndrome Scale (t-score) |
| Fluid Intelligence | ABCD NIH Toolbox: Fluid Composite | abcd_tbss01: nihtbx_fluidcomp_agecorrected | Age-Corrected Standard Score |
| Crystalised Intelligence | ABCD NIH Toolbox: Crystallized Composite | abcd_tbss01: nihtbx_cryst_agecorrected | Age-Corrected Standard Score |
| **Academic Performance and School Engagement** | | | |
| Drop in grades | ABCD Parent Diagnostic Interview for DSM-5 Background Items; Parents answered the following question: In the past year or past several months, has there been a drop in child's grades? | dibf01: kbi_p_c_drop_in_grades | 1 = Yes; 2 = No |
| In-School Special Services: Emotional Support | ABCD Parent Diagnostic Interview for DSM-5 Background Items Question; Parents answered the following question: “Does your child receive special services at school? Full-time Emotional Support Classroom | dibf01: kbi_p_c_spec_serv___1 | 1 = Yes; 2 = No |
| In-School Special Services: Learning Support | ABCD Parent Diagnostic Interview for DSM-5 Background Items; Parents answered the following question: ““Does your child receive special services at school? Full-time Learning Support Classroom | dibf01: kbi_p_c_spec_serv___2 | 1 = Yes; 2 = No |
| In-School Aide | ABCD Parent Diagnostic Interview for DSM-5 Background Items. Parents answered the following question: “Does your child receive special services at school? Full-time or Part-time Aide | dibf01:  kbi_p_c_spec_serv___3 (full-time)  kbi_p_c_spec_serv___5 (part-time) | 1 = has either part-time or full-time aide  0 = has no aide |
| School disengagement  Child Report | ABCD School Risk and Protective Factors Survey (SRPF)- Disengagement Subscale; Two questions reflecting alienation from academic goals | abcd_sscey01: srpf_y_ss_dfs | Responses to each question were coded as:  1=Strongly Disagree  2=Disagree  3=Agree  4=Strongly Disagree  The sum of the responses was used; higher values denote greater disengagement |
| School involvement  Child Report | ABCD School Risk and Protective Factors Survey - Involvement Subscale; Four questions reflecting positive involvement at school | abcd_sscey01; srpf_y_ss_iiss | Responses to each question were coded as:  1=Strongly Disagree  2=Disagree  3=Agree  4=Strongly Disagree  The sum of the responses was used; higher values denote greater engagement in academic activities |
| Positive View of School Environment  Child report | ABCD School Risk and Protective Factors Survey - Environment Subscale; Six questions that tap into the child’s experience of school as an environment of opportunity and support | abcd_sscey01: srpf_y_ss_ses | Responses to each question were coded as:  1=Strongly Disagree  2=Disagree  3=Agree  4=Strongly Disagree  The sum of the responses was used; higher values denote a more positive view of the school environment |
| **Parental Characteristics** | | | |
| Parental education  Parent report | ABCD Parent Demographics Survey; Participants answered the following question: “What is the highest grade or level of school you have completed or the highest degree you have received?” | pdem02: demo_prnt_ed_v2 | 1=1st grade  2=2nd grade  3=3rd grade  4=4th grade  5=5th grade  6=6th grade  7=7th grade  8=8th grade  9=9th grade  10=10th grade  11=11th grade  12=12th grade  13=High School  14= GED or equivalent diploma  15=Some college  16=Associate degree: occupational  17=Associate degree: Academic program  18=Bachelor’s degree  19=Master’s degree  20=Professional School degree  21=Doctoral degree |
| Parental employment  Parent report | ABCD Parent Demographics Survey; Participants answered the following question: “Are you working now, looking for work, retired, stay at home parent, a student, or something else?” | pdem02: demo_prnt_empl_v2 | 1=currently employed full time/part time;  0=if endorsed any of the following:  temporarily Laid off, looking for work,  retired, disabled, stay at home, student,  sick Leave, maternity Leave,  unemployed not looking for work |
| Parental partnership  Parent report | ABCD Parent Demographics Survey; Parents answered the following question: “Are you now married, widowed, divorced, separated, never married or living with a partner” | pdem02: demo_prnt_marital_v2 | Responses are coded as:  1=married/living with partner  0=widowed/divorced/separated/never married |
| Parental Psychiatric Diagnosis  Parent report | ABCD Parent Family History Summary Scores; this instrument provides summary scores from the family history inventory that assesses the lifetime occurrences of a range of psychological problems in all first- and second-degree, biological relatives of the child, as reported by a caregiver. | abcd_fhxssp01:  famhx_ss_parent_alc_p  famhx_ss_parent_dg_p  famhx_ss_parent_dprs_p  famhx_ss_parent_ma_p  famhx_ss_parent_vs_p  famhx_ss_parent_trb_p  famhx_ss_parent_nrv_p  famhx_ss_parent_hspd_p  famhx_ss_parent_scd_p | Responses for each diagnosis listed below were coded as:  0=neither parent; 1=one parent; 2=both parents  Alcohol Related Diagnoses famhx_ss_parent_alc_p  Drug Related Diagnoses famhx_ss_parent_dg_p  Depressive Disorders famhx_ss_parent_dprs_p  Mania  famhx_ss_parent_ma_p  Paranoia  famhx_ss_parent_vs_p  Antisocial Behavior  famhx_ss_parent_trb_p  Nervous breakdown famhx_ss_parent_nrv_p  Psychiatric hospitalization famhx_ss_parent_hspd_p  Attempted/committed suicide famhx_ss_parent_scd_p |
| Parental Help Seeking for Mental Problems  Parent report | ABCD Parent Family History Summary Scores; this instrument provides summary scores from the family history inventory that assesses the lifetime occurrences of a range of psychological problems in all first- and second-degree, biological relatives of the child, as reported by a caregiver. | abcd_fhxssp01:  famhx_ss_parent_prf_p | Seen a professional for overall mental/emotional problem  famhx_ss_parent_prf_p |
| **Family and Peer Environment** | | | |
| Family income  Parent report | ABCD Parent Demographics Survey; Participants answered the following question: “What is your total combined family income for the past 12 months?” | pdem02: demo_comb_income_v2 | 1= Less than $5,000;  2=$5,000 through $11,999;  3=$12,000 through $15,999;  4=$16,000 through $24,999;  5=$25,000 through $34,999; 6=$35,000 through $49,999;  7=$50,000 through $74,999;  8= $75,000 through $99,999;  9=$100,000 through $199,999;  10=$200,000 and greater. |
| Family financial adversity  Parent report | ABCD Parent Demographics Survey; | pdem02:  demo_fam_exp1_v2  demo_fam_exp2_v2 demo_fam_exp3_v2  demo_fam_exp4_v2  demo_fam_exp5_v2 demo_fam_exp6_v2 demo_fam_exp7_v2 | demo_fam_exp1_v2: Needed food but couldn't afford to buy it or couldn't afford to go out to get it  demo_fam_exp2_v2: Without telephone service because could not afford it  demo_fam_exp3_v2: Didn't pay the full amount of the rent or mortgage because could not afford it  demo_fam_exp4_v2: Evicted from home for not paying the rent or mortgage  demo_fam_exp5_v2: Had services turned off by the gas or electric company, or the oil company wouldn't deliver oil because payments were not made  demo_fam_exp6_v2: Had someone who needed to see a doctor or go to the hospital but didn't go because could not afford it  demo_fam_exp7_v2: Had someone who needed a dentist but couldn't go because could not afford it  The sum of all responses is used; higher values denote greater financial adversity. |
| Family conflict  Parent report | ABCD Family Environment Scale: Family Conflict Subscale; Modified from PhenX | abcd_sscep01: fes_p_ss_fc  fes_enviro1_q1 + fes_enviro1_q2 + fes_enviro1_q3 + fes_enviro1_q4 + fes_enviro1_q5 + fes_enviro1_q6 + fes_enviro1_q7 + fes_enviro1_q8 + fes_enviro1_q9 | The number of times events indicating family conflict occurred in the past year are summed for all questions; higher scores indicating greater conflict. |
| Family conflict  Child report | ABCD Family Environment Scale: Family Conflict Subscale; Modified from PhenX | abcd_sscey01: fes_y_ss_fc  fes_youth_q1 + fes_youth_q2 + fes_youth_q3 + fes_youth_q4 + fes_youth_q5 + fes_youth_q6 + fes_youth_q7 + fes_youth_q8 + fes_youth_q9 | The number of times events indicating family conflict occurred in the past year are summed for all questions; higher scores indicating greater conflict. |
| Primary caregiver's warmth  Child report | Acceptance subscale from the Children's Report of Parental Behavioral Inventory; 5 questions: respond to items describing the primary caregivers’ behaviors that indicate warmth or acceptance | abcd_sscey01: crpbi_y_ss_parent | Responses to each item were coded as:  0=No  2=Somewhat  3=A lot  The mean of the responses was used; higher values denote greater caregiver warmth |
| Parental supervision  Child report |  | abcd_sscey01: pmq_y_ss_mean | Responses to each item were coded as:  1=Never  2=Almost never  3=Sometimes  4=Often  5=Always or almost always  The mean of the responses was used; higher values denote more parental supervision |
| Child has a regular group of friends  Parent report | Does the child have a regular group of kids he or she hangs out with at school or in your neighborhood? | dibf01: kbi_p_c_reg_friend_group | 1 = Yes; 0= No |
| Bullying  Parent report | Does the child have any problems with bullying at school or in your neighborhood. | dibf01: kbi_p_c_bully | 1=Yes; 0=No. |
| Exposure to stressful events  Parent report | KSADS-5 Section on Traumatic Events | abcd_ptsd01:  ksads_ptsd_raw_754_p  ksads_ptsd_raw_755_p  ksads_ptsd_raw_756_p  ksads_ptsd_raw_757_p  ksads_ptsd_raw_758_p  ksads_ptsd_raw_759_p  ksads_ptsd_raw_760_p  ksads_ptsd_raw_761_p  ksads_ptsd_raw_762_p  ksads_ptsd_raw_763_p  ksads_ptsd_raw_764_p  ksads_ptsd_raw_765_p  ksads_ptsd_raw_766_p  ksads_ptsd_raw_767_p  ksads_ptsd_raw_768_p  ksads_ptsd_raw_769_p  ksads_ptsd_raw_770_p | Responses to each stressful event were coded as:  0=Never exposed  1=Exposed  ksads_ptsd_raw_754_p: A car accident in which your child or another person in the car was hurt bad enough to require medical attention  ksads_ptsd_raw_755_p: Another significant accident for which your child needed specialized and intensive medical treatment  ksads_ptsd_raw_756_p: Witnessed or caught in a fire that caused significant property damage or personal injury  ksads_ptsd_raw_757_p: Witnessed or caught in a natural disaster that caused significant property damage or personal injury  ksads_ptsd_raw_758_p: Witnessed or present during an act of terrorism  ksads_ptsd_raw_759_p: Witnessed death or mass destruction in a war zone  ksads_ptsd_raw_760_p: Witnessed someone shot or stabbed in the community  ksads_ptsd_raw_761_p: Shot, stabbed, or beaten brutally by a non-family member  ksads_ptsd_raw_762_p: Shot, stabbed, or beaten brutally by a grown up in the home  ksads_ptsd_raw_763_p: Beaten to the point of having bruises by a grown up in the home  ksads_ptsd_raw_764_p: A non-family member threatened to kill your child  ksads_ptsd_raw_765_p: A family member threatened to kill your child  ksads_ptsd_raw_766_p: Witness the grownups in the home push, shove or hit one another  ksads_ptsd_raw_767_p: A grown up in the home touched your child in their privates, had your child touch their privates, or did other sexual things to your child  ksads_ptsd_raw_768_p: An adult outside your family touched your child in their privates, had your child touch their privates or did other sexual things to your child  ksads_ptsd_raw_769_p: A peer forced your child to do something sexually  ksads_ptsd_raw_770_p: Learned about the sudden unexpected death of a loved one  1=endorsing any of the above adverse events  0=endorsing none of the above adverse events |
| **Neighbourhood Environment** | | | |
| Quality of Neighbourhood Environment | Area Deprivation Index (ADI)  Derived from Residential History | abcd_rhds01: reshist_addr1_adi_perc | ADI national percentiles; higher means higher value of ADI |
| Caregiver refers to the individual completing the parental portion of the instruments | | | |

**S5. Neuroimaging** **acquisition, processing and feature extraction**

Neuroimaging data from the ABCD participants were acquired on 3T scanners (Siemens Prisma, General Electric MR 750, Philips) using standardized sequences [7]. The study also employed standardized quality assurance and processing protocols [8].

Brain Morphometry: 3D T1-weighted scans were acquired using magnetization-prepared rapid acquisition gradient echo sequence: matrix = 256×256; voxel resolution = 1×1×1 mm3 isotropic; repetition time (TR) = 2500 ms; echo time (TE) = 2.88 ms; field of view (FOV) = 256×256 mm; flip angle = 8 degrees with prospective motion correction. These images were processed using standard pipelines implemented in the FreeSurfer image analysis suite (version 5.3.0; http://surfer.nmr.mgh.harvard.edu/) for cortical reconstruction (based on the Desikan-Killiany Atlas [9]) and volumetric segmentation of subcortical structures (based on the probabilistic atlas in FreeSurfer [10]. This procedure yielded 68 cortical thickness measures, 68 cortical area measures, and 16 subcortical volumetric measures.

Intracortical Myelination: An estimate of myelination was obtained through the gray/white matter contrast (GWC) from the T1-weighted images. Intensity values from the T1-weighted images of the ABCD participants were sampled at a distance of ±0.2 mm - relative to the gray-white boundary along the normal vector at each surface location; the GWC was computed as ([white - gray]/[white + gray]/2) and regional averages were then computed for each Desikan-Killiany atlas defined cortical region.

Neurite Density: Diffusion MRI (dMRI) data were acquired using the following parameters: multiband EPI with slice acceleration factor 3 and 96 diffusion directions, 1.7 mm isotropic, seven b = 0 frames, and four b-values (6 directions with b = 500 s/mm2, 15 directions with b = 1000 s/mm2, 15 directions with b = 2000 s/mm2, and 60 directions with b = 3000 s/mm2). Restriction Spectrum Imaging (RSI) [11] was applied to the dMRI data to separate intracellular restricted diffusion from hindered diffusion across different length scales and tissue geometries to yield information on tissue cellularity. The volume fraction of anisotropic restricted diffusion (the shortest length scale examined) is believed to reflect the relative density of neuronal processes (neurite density, ND). RSI data were corrected for motion and eddy current distortions, spatial and intensity distortions, and distortions caused by gradient nonlinearities, before estimating the ND. Mean ND values were generated for each subcortical and cortical region (as defined using the same parcellation as for the morphological data).

**S6. Heterogeneity Through Discriminative Analysis (HYDRA)**

**Supplementary Figure S2. Schematic representation of HYDRA.** The model uses multiple classifiers that form linear hyperplanes (black lines) whose segments separate cases from controls into n clusters by the largest margin. The number of clusters is determined empirically using the Adjusted Rand Index [12].

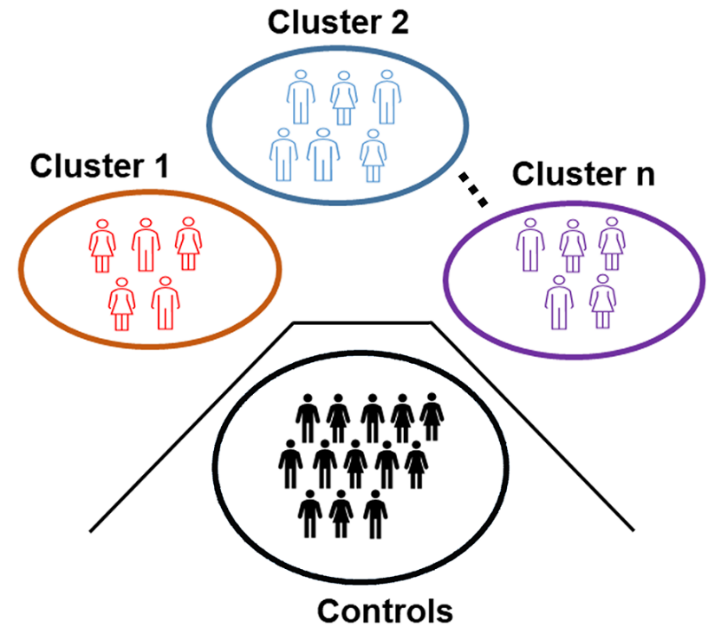

**Permutation Analysis:** We conducted permutation tests to assess the stability of the observed Adjusted Rand index (ARI) [12] to changes in the sample composition following published procedures [13, 14]. First, we assessed whether the solution obtained in the main analyses differed from that generated following random grouping of participants. To achieve this, we compared the ARI observed in the main analyses to that generated by HYDRA after randomly shuffling the group labels of typical-developing participants and participants with mood and anxiety disorders while maintaining the control-to-patient ratio of the original data and permuting 100 times. Second, we assessed whether the solution obtained in the main analyses differed from that generated if the sample did not include a clinical group (i.e., if disorder-related variability was removed). To achieve this, we compared the ARI observed in the main analyses to that generated by HYDRA after randomly assigning typically developing participants (n=2823) to a “control group” and to a “pseudo-patient group” while maintaining the control-to-patient ratio of the original study (i.e., controls n=1676 and pseudo-patients n=1147) and permuting 100 times.

**Supplementary Figure S3**. **Correlations between total intracranial volume (TIV) and global cortical thickness, surface area, and subcortical volume.** The relationships were fitted between TIV and cortical thickness averaged across both hemispheres (left panel), between TIV and total cortical surface area (middle panel), and between TIV and total subcortical volume (right panel). Significant linear effects were observed for surface area and subcortical volume.

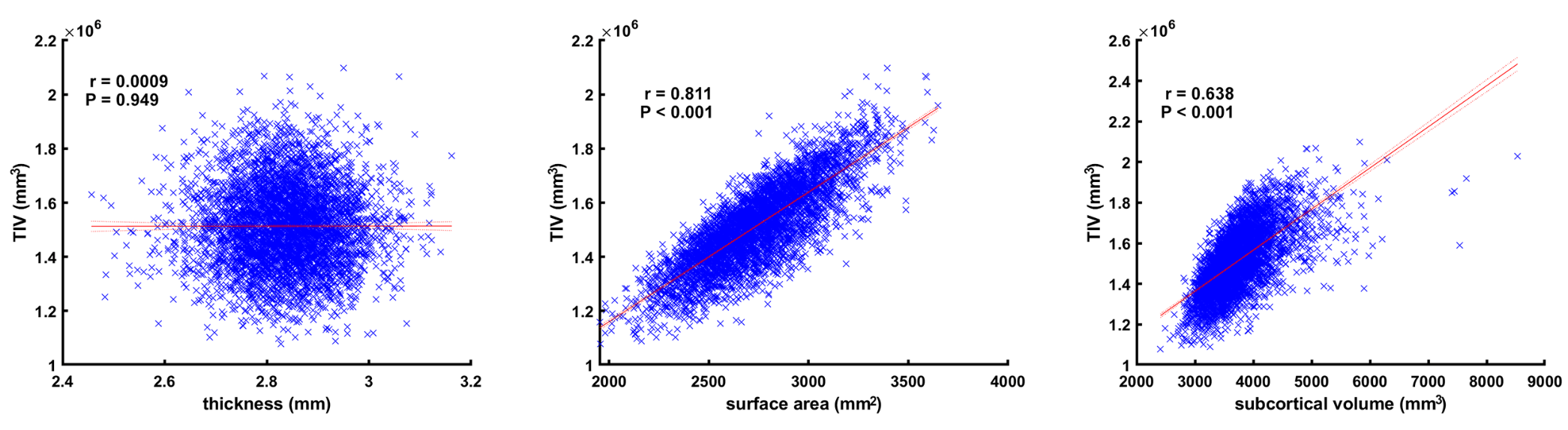

**Supplementary Figure S4**. **Correlations between total intracranial volume (TIV) and global gray/white matter contrast (GWC) and neurite density.** The relationships were fitted between TIV and GWC averaged across both hemispheres (left panel), between TIV and cortical neurite density (middle panel), and between TIV and subcortical neurite density (right panel). Significant linear effects were observed for GWC and cortical neurite density.

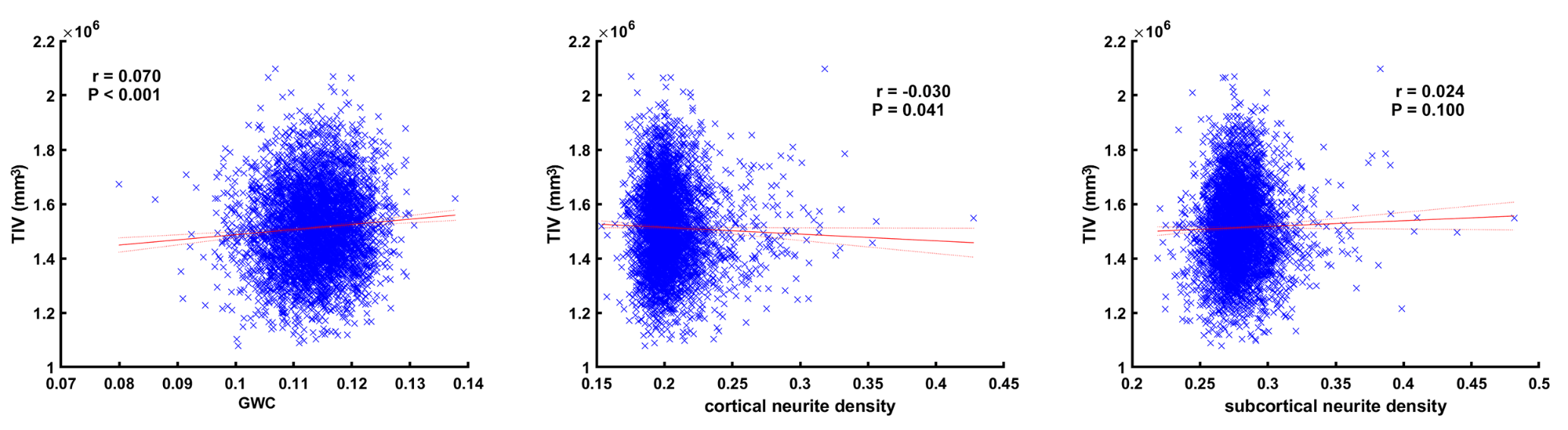

**S**

**S7. Sensitivity Analyses**

**Reproducibility:** Inter-study reproducibility by splitting the sample into a discovery subset (80%) and a reproducibility subset (20%) with each subset retaining the demographic and diagnostic distribution of the whole sample. The HYDRA was applied to the two subsets following the procedures outlined for the main analyses and to test the robustness of the ARI across the original sample, and the discovery and reproducibility subsets.

**Intracortical Myelin:** The T_1w_-T_2w_ ratio is another measure thought to be sensitive to intracortical myelination [15]. We used Pearson correlation to test the spatial similarity between the GWC and T1w/T2w cortical maps of each of the subtypes identified by HYDRA in the main analyses.

**Relatedness:** We randomly excluded siblings from the same families so that in the relatedness subsample only one member per family was included. This procedure resulted in the exclusion of 4.9% of the original study sample. We then repeated the statistical analyses. There were no siblings amongst participants diagnosed with depressive or comorbid anxiety and mood disorders. The percentage of excluded participants in the typically developing group, the bipolar disorders group, and the anxiety disorders group was 7.16%, 1.30%, and 2.59% respectively.

**Supplementary Results**

**S8. HYDRA identified 3 subtypes of youth with mood and anxiety disorders**

Of the six clustering solutions considered (from 2 to 7), a well-defined peak at *K* = 3 emerged, suggesting the presence of three distinct subtypes of participants with mood and anxiety disorders. The observed ARI for the 3-cluster solution was statistically significantly higher in the observed ARI compared to the first null distribution (P=0.01). ARIs for all other observed clustering solutions (i.e., 3, 4, 5, 6, and 7) showed no significant differences to the first null distribution (all P>0.50). Similarly, the observed ARI for the 3-cluster solution was significantly higher than the second null distribution (P=0.05), whereas ARIs for all other observed clustering solutions showed no significant differences to the second null distribution (all P>0.50). These results confirmed the 3-cluster solution as the optimal clustering solution.

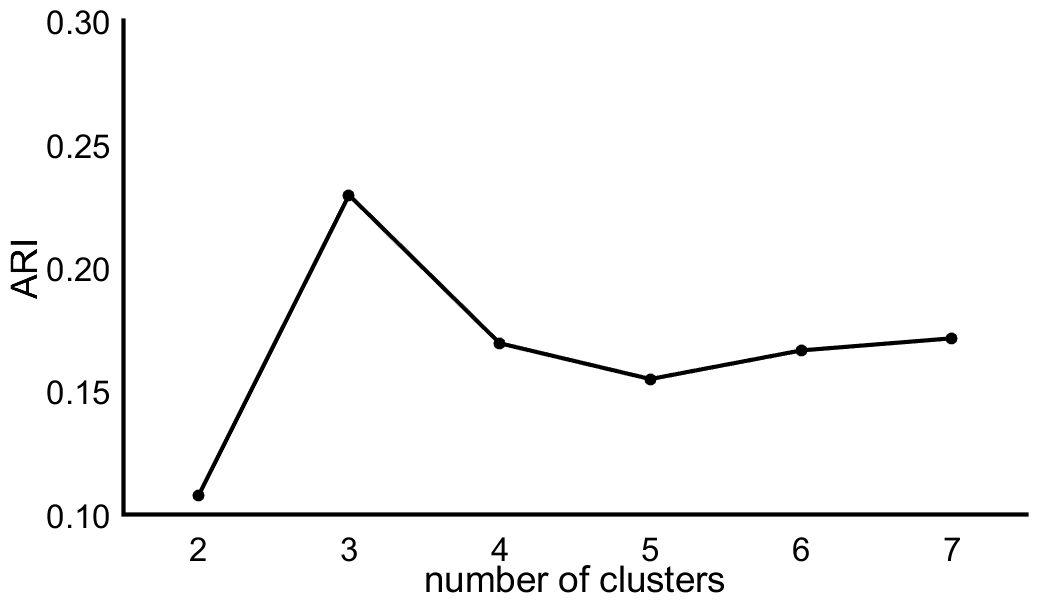

**Supplementary Figure S5. Adjusted Rand Index (ARI) for HYDRA Models of different cluster solutions.**

| Supplementary Table S5. Clinical, family, and social characteristics of the HYDRA subtypes that did not differ from the typically developing group | | | | | |
| --- | --- | --- | --- | --- | --- |
|  | **Typically**  **Developing**  **N=2823** | | **Subtype 1**  **N=631** | **Subtype 2**  **N=649** | **Subtype 3**  **N=651** |
| Obstetric and Perinatal | | | | | |
| Number of Obstetric Complications, mean (SD) | | 0.32 (0.68) | 0.39 (0.76) | 0.35 (0.7) | 0.35 (0.79) |
| Premature birth (%yes/%no) | | 19/81 | 19/81 | 17/83 | 20/80 |
| Number of maternal medical conditions during pregnancy, mean (SD) | | 0.85 (2.78) | 1.07 (3.06) | 0.98 (2.93) | 1.09 (3.03) |
| Academic Performance and School Environment | | | | | |
| In-school Emotional Support (%yes/%no) | | 0.2/99.8 | 0.3/99.7 | 0.3/99.7 | 0.5/99.5 |
| In-School Learning Support (%yes/%no) | | 1/99 | 0.3/99.7 | 1/99 | 1/99 |
| In-School Aid (%yes/%no) | | 1/99 | 0.6/99.4 | 2/98 | 1/99 |
| Parental Characteristics | | | | | |
| Parental Employment (%yes/%no) | | 72/28 | 71/29 | 70/30 | 69/31 |
| Family and Peer Environment | | | | | |
| Primary Caregiver’s Warmth, mean (SD) | | 2.81 (0.27) | 2.79 (0.28) | 2.79 (0.29) | 2.8 (0.28) |
| Having a Best Friend (%yes/%no) | | 85/15 | 85/15 | 87/13 | 86/14 |
| Continuous variables are shown as mean (standard deviation; SD) and categorical variables as percentage (%). Definitions of all the variables are provided in Supplemental Table 4. Variables showing significant group differences at P_FDR_<0.005 are detailed in Table 1 in the main manuscript | | | | | |

**S9. Sensitivity Analysis**

**Reproducibility:** The reproducibility analyses confirmed the robustness of a *K*=3 solution as optimal.

**Intracortical Myelin:** The correlation between cortical maps for the GWC and the T_1w_/T_2w_ ratio within each HYDRA subtype and within the typically developing group was highly correlated suggesting that both measures are similarly informative. Figure S6 depicts the mean GWC surface map of each subtype compared to the mean T1w/T2w ratio map of the corresponding subtype. The absolute correlation coefficients of the two mean maps for the typically developing group and for subtype 1, subtype 2, and subtype 3 were 0.70, 0.69, 0.70, and 0.69 respectively.

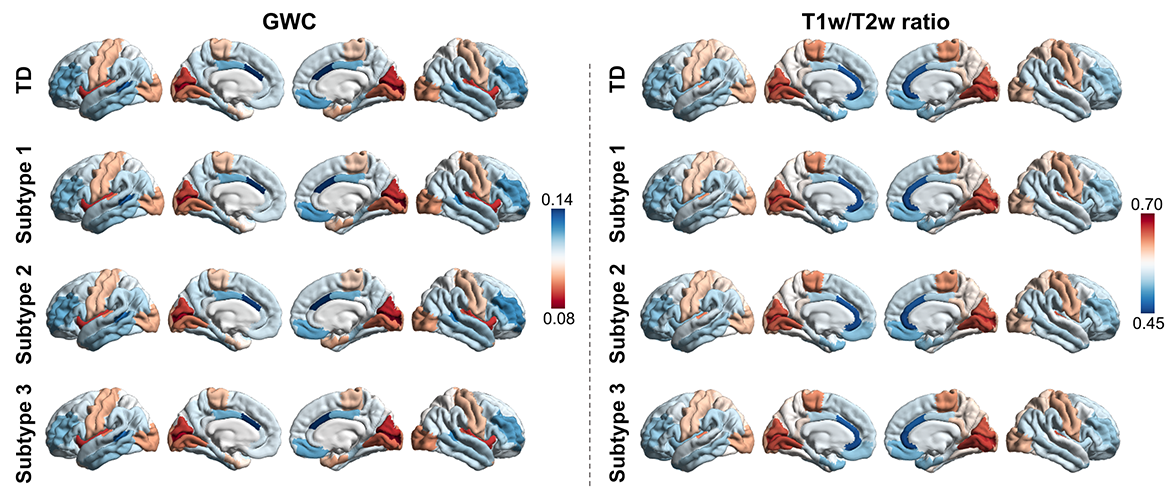

**Supplementary Figure S6. Cortical maps for GWC and T1w/T2w ratio within each HYDRA subtype and typically developing group (TD).** For visualization purposes, for the GWC maps, warm colors represent regions with lower gray/white matter contrast, while cold colors represent regions with higher gray/white matter contrast; for the T1w/T2w ratio map, warm colors are thought to represent regions with a high amount of intracortical myelin, while cold colors represent regions of low myelin content.

**Relatedness:** Exclusion of related individuals from the analyses did not alter the results of the main analyses as illustrated in Supplementary Figure S7. The correlation coefficients between the t-valued spatial maps of the main results and that of the sensitivity analysis were all r>0.99.

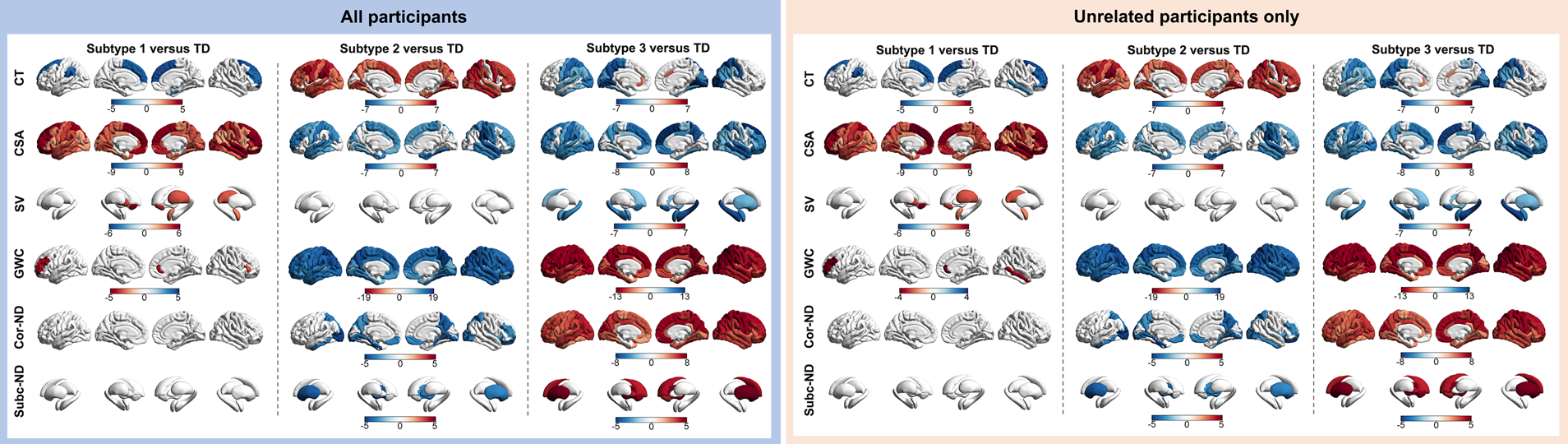

**Supplementary Figure S7. Neuroimaging Profiles of the HYDRA-identified Subtypes in the entire sample (left panel) and after exclusion of all related individuals (right panel).** Regional differences (represented by the corresponding t-values) between each subtype and the typically developing (TD) group in cortical thickness (CT), cortical surface area (CSA), subcortical volumes (SV), neurite density (ND) and gray/white matter contrast (GWC). Only significant differences at PFDR<0.005 are shown. Warm and cool colors respectively indicate higher and lower values in each subtype compared to the TD group; however, for visualization purposes, in the GWC plots warm and cool colors respectively indicate lower and higher values in each subtype compared to the TD group.
